# Ripple Band Phase Precession of Place Cell Firing during Replay

**DOI:** 10.1101/2021.04.05.438482

**Authors:** Daniel Bush, Freyja Olafsdottir, Caswell Barry, Neil Burgess

## Abstract

Phase coding offers several theoretical advantages for information transmission compared to an equivalent rate code. Phase coding is shown by place cells in the rodent hippocampal formation, which fire at progressively earlier phases of the movement related 6-12Hz theta rhythm as their spatial receptive fields are traversed. Importantly, however, phase coding is independent of carrier frequency, and so we asked whether it might also be exhibited by place cells during 150-250Hz ripple band activity, when they are thought to replay information to neocortex. We demonstrate that place cells which fire multiple spikes during candidate replay events do so at progressively earlier ripple phases, and that spikes fired across all replay events exhibit a negative relationship between decoded location within the firing field and ripple phase. These results provide insights into the mechanisms underlying phase coding and place cell replay, as well as the neural code propagated to downstream neurons.

## Introduction

The mammalian hippocampus is implicated in both spatial cognition and episodic memory function (O’Keefe and Nadel, 1978; Eichenbaum, 2000). In the rodent hippocampal formation, place and grid cells are active in restricted regions of space – the corresponding place or grid field (O’Keefe and Dostrovsky, 1971; Hafting et al., 2005). During active movement, when 6-12Hz theta oscillations dominate the local field potential (LFP), place cells and a subset of grid cells in medial entorhinal cortex (MEC) also exhibit a theta phase code for location. Specifically, these cells fire at progressively earlier phases of the theta cycle as the firing field is traversed (O’Keefe and Recce, 1993; Hafting et al., 2008). Because phase precession is coordinated across cells, this produces theta ‘sweeps’ of activity at the network level that encode a sequence of locations beginning behind and progressing ahead of the animal within each oscillatory cycle (Burgess et al., 1994; Skaggs et al., 1996; Johnson and Redish, 2007). The theta phase code for location exhibited by place and grid cells improves the accuracy of decoding location (Jensen and Lisman, 2000) and allows movement direction to be inferred from population activity in each oscillatory cycle (Zutshi et al., 2017; Bush and Burgess, 2020).

Importantly, however, phase is independent of frequency, and a similar coding scheme could therefore be supported by oscillatory activity in other frequency bands, such as 25-55Hz slow gamma (Fries et al., 2007; Zheng et al., 2016); or by an LFP signal whose instantaneous frequency varies dynamically over a wide range, such as that observed in bats and humans (Jacobs, 2014; Eliav et al., 2018; Bush and Burgess, 2020; Qasim et al., 2020). During periods of quiescent waking and rest, the hippocampal LFP exhibits prominent sharp-wave ripple (SWR) events comprised of a large amplitude deflection accompanied by a transient increase in 150-250Hz ripple band power (Buzsaki et al., 1992). SWR events are associated with prominent place cell multi-unit activity (MUA), which can recapitulate coherent spatial trajectories through recently visited environments (Wilson and McNaughton, 1994; O’Neill et al., 2010; Olafsdottir et al., 2018). Here, we asked whether place cells might also exhibit phase coding relative to ripple band oscillations during replay events. Specifically, this should be characterised by a systematic change in ripple band firing phase across multiple spikes fired by individual place cells within each candidate replay event; a relationship between decoded location within the firing field and ripple band firing phase across multiple replay events through the same place field; and a relationship between ripple band phase and the relative location encoded by population activity across multiple place cells within each replay event.

## Results

We analysed data from rats completing shuttle runs along a linear track for food reward and during subsequent sleep (as described previously: Olafsdottir et al., 2016; 2017). Six animals undertook a total of 29 RUN sessions on different days lasting 34.4±11.7 minutes (median±SD, range 21.6–60.9). During RUN sessions, animals successfully completed 20±5.7 outbound and inbound runs along a 6m Z-shaped track (Figure 1A) while we recorded the activity of 34±17.7 putative pyramidal cells in dorsal CA1 (14–71 per session, 1044 in total). After each RUN session, animals rested for 93.5±10.1 (90-129) minutes (REST) while we continued recording the activity of 30±17.8 putative pyramidal cells in dorsal CA1 (14–76 per session, 1025 in total, including 960 that were also active during RUN) alongside local field potential (LFP) at a high sample rate (4.8kHz). Putative interneurons, identified by narrow waveforms and mean firing rates >10Hz, were excluded from all analyses.

**Figure 1:**
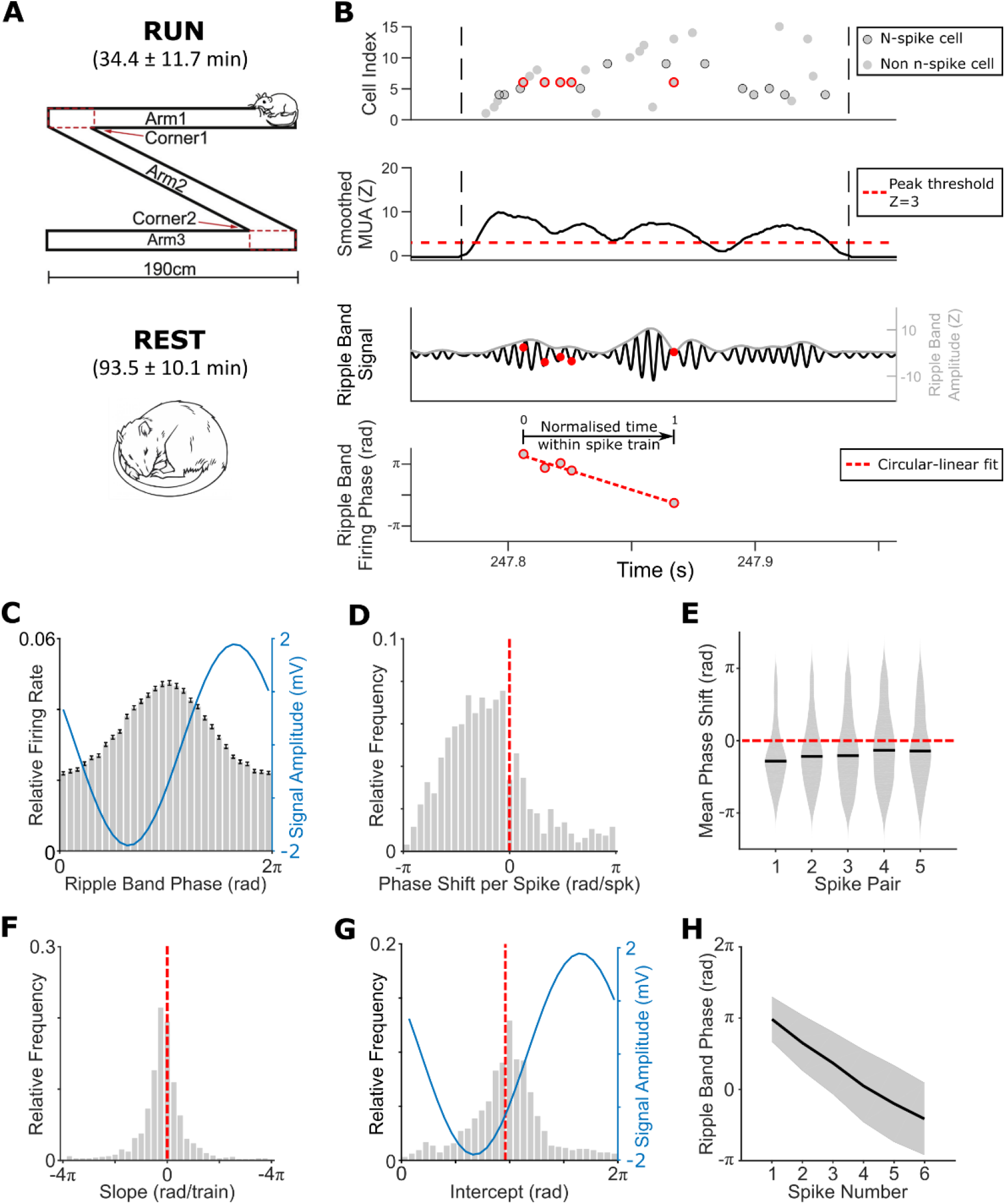
Changes in place cell ripple band firing phase during candidate replay events. **[A]** Task schematic; **[B]** Schematic analysis of an ‘n-spike’ event, spikes highlighted in red correspond to the multiple spikes fired by a single ‘n-spike’ cell, and plotted in the bottom panels; **[C]** Relative firing rate (grey bars) and mean signal amplitude (blue line) by ripple band phase; **[D]** Circular mean phase shift between all successive spikes in candidate replay events for each n-spike cell (n=953, overall circular median±circular SD=-0.922±1.19 rad/spk). This distribution is non-uniform (Rayleigh test, z=229, p<0.001) with a median value that differs significantly from zero (circular median test, p<0.001); **[E]** Circular mean phase shift by within-event spike pair, averaged across all cells. Each phase shift is non-uniformly distributed with a median value that differs significantly from zero (all p<0.001); **[F]** Distribution of normalised time within spike train vs. ripple band phase slopes across cells (overall median±SD=-0.502±2.66 rad), which differs significantly from zero (t(949)=-6.27, p<0.001); **[G]** Distribution of normalised time within spike train vs. ripple band phase intercepts across cells (overall circular mean±circular SD=3.06±1.04 rad). This distribution is non-uniform (Rayleigh test, z=323, p<0.001) with a median value that differs significantly from the preferred firing phase of each cell (3.01±0.79 rad, indicated by the dashed red line; circular median test, p<0.01); **[H]** Unwrapped ripple band phase by spike number within candidate replay events, averaged across all cells (error bars indicate circular standard deviation)

First, we looked for candidate replay events during REST on the basis of multi-unit activity (MUA; following Olafsdottir et al., 2017, see Methods for further details). This identified a total of 25328 events (758±599 per session, range 97–3113, equivalent to events occurring a rate of ∼0.13Hz). Candidate events lasted 114±71.3ms and incorporated activity in 23.3±10.2% of all recorded pyramidal cells (range 15.0-86.2%; see Figure 1B for an example, Figure S1 for further details). During these candidate replay events, average power in the 150-250Hz ripple band was elevated (mean Z=1.98±1.52, peak Z=7.17±4.36, peak frequency=194±11.6Hz), and the firing rate of 69.4% of pyramidal cells was significantly modulated by ripple band phase (Figure 1C). Although active cells typically fired only a single spike each during candidate replay events (median±SD=1±1.4), multiple cells were active during each event. Hence, in order to examine changes in the ripple band firing phase of individual cells over the course of each event, we identified a subset of ‘n-spike’ events during which one or more active cells fired ≥3 spikes each. The vast majority of recorded cells (93.0%) participated in at least one n-spike event, the majority of candidate replay events (20224, or 79.9%) were n-spike events (range 60.3–94.0% per session), and n-spike events tended to be longer (127ms vs. 71ms; Mann-Whitney U-test, Z=70, p<0.001) and incorporate more active cells (24.1% vs. 20.7%; Mann-Whitney U-test, Z=27.8, p<0.001) than other events.

Next, we estimated the ripple band firing phase of each spike by applying the Hilbert transform to LFP data filtered in the 150-250Hz range - shifting the resultant phase values so that π rad corresponded to the circular mean firing phase of all cells recorded in each session. We then computed the ripple band phase shift between successive pairs of spikes fired by n-spike cells, averaged those phase shifts across all spike pairs, and then across all n-spike events. Remarkably, we found that these phase shifts were consistently negative (Figure 1D). Specifically, the distribution of circular mean phase shifts across cells was non-uniform, with an overall circular median that differed significantly from zero. Importantly, this was true when each spike pair (i.e. first, second, third etc.) was considered separately, indicating that the overall effect was consistent across the spike train (Figure 1E). In addition, we used circular-linear regression to estimate the slope and intercept of the relationship between normalised time within the spike train (i.e. where the time of the first / last spike was 0 / 1, respectively) and ripple band phase for each n-spike cell in each n-spike event (Kempter et al., 2012; see Methods for further details). Consistent with the results above, the median slope of this relationship was negative and significantly different from zero across cells (Figure 1F). Importantly, the distribution of intercepts was also non-uniform, with a circular mean value that was slightly but significantly later in the ripple cycle than the mean firing phase of all n-spike cells (Figure 1G), consistent with a shift to earlier phases when spikes occur later in the train. Hence, the ripple band firing phase of putative pyramidal cells during candidate replay events begins just after the phase of peak firing and becomes progressively earlier as the event continues (Figure 1H).

Importantly, when analysed separately, each animal exhibited negative circular mean phase shifts (range of -1.57 to -0.43 rad/spk; circular median test, p<0.05) and mean slopes (range of -0.657 to - 0.394 rad, t(5)=-15.1, p<0.001) averaged across all n-spike cells. Moreover, because n-spike cells typically fire <1 spike per ripple band cycle (0.179±0.108 across all events, Figure S1I), results were similar if we included only the first spike from each oscillatory cycle in these analyses (see Figure S2 for details). Qualitatively similar results were also obtained when analysing candidate ‘online’ replay events, which occurred while the animal was awake but immobile at the corners of the track during RUN, indicating that this phenomenon is preserved across behavioural states (see Figure S3 for details); and when candidate replay events were identified using ripple band power instead of MUA (see Figure S4, Methods for details). Conversely, we observed no consistent change in ripple band firing phase across n-spike trains fired by putative principal neurons recorded simultaneously in the deeper layers of medial entorhinal cortex (MEC), nor if we restricted that analysis to just grid cells (see Figure S5 for details).

Next, for comparison, we used analogous methods to examine changes in theta firing phase across spike trains from the same cells during active movement on the track (Figure 2A). Specifically, we looked for extended periods of elevated activity in each cell, independent of spatial location, and inspected the relationship between theta firing phase – again, defined such that π rad corresponds to the circular mean preferred firing phase of all cells recorded in each session - and spike number or normalised time within each n-spike train (Aghajan et al., 2014; see Methods for further details, Figure S6 for details of theta n-spike trains). During these theta n-spike trains, the firing rate of 37.1% of pyramidal cells was significantly modulated by theta phase (Figure 2B). In this case, because active cells typically fire >1 spike per theta cycle (1.57±3.0 across all cells in all n-spike trains), we restricted our subsequent analyses to the first spike from each oscillatory cycle (see Methods).

**Figure 2:**
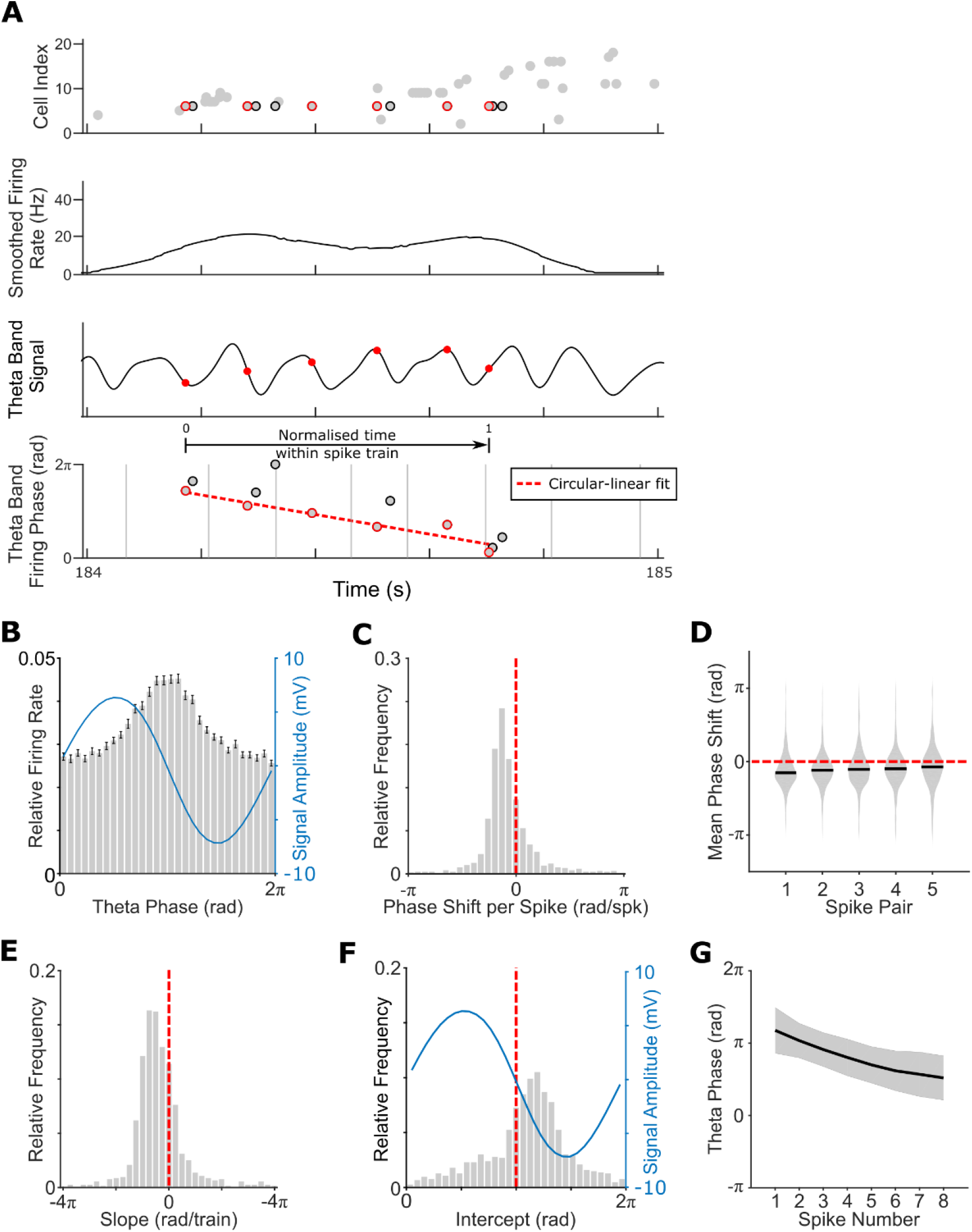
Changes in place cell theta firing phase during n-spike trains. **[A]** Schematic analysis of a theta ‘n-spike train’, spikes highlighted in black correspond to those fired by a single illustrative cell and plotted in the lower panels, spikes highlighted in red correspond to the first in each theta cycle; **[B]** Relative firing rate (grey bars) and mean signal amplitude (blue line) by theta phase; **[C]** Circular mean phase shift between successive spikes in theta n-spike trains (overall circular mean±circular SD=-0.307±0.61 rad/spk). This distribution is non-uniform (Rayleigh test, z=673, p<0.001) with a median value that differs significantly from zero (circular median test, p<0.001); **[D]** Circular mean phase shift by within-event spike pair, each of which are non-uniformly distributed with a median value that differs significantly from zero (all p<0.001); **[E]** Distribution of normalised within n-spike train time vs. theta phase slopes (overall median±SD=-1.44±2.77 rad), which differs significantly from zero (t(977)=-13.1, p<0.001); **[F]** Distribution of normalised within n-spike train time vs. theta phase intercepts across cells (overall circular mean±circular SD=3.59±1.12 rad). This distribution is non-uniform (Rayleigh test, z=285, p<0.001) with a median value that differs significantly from the preferred firing phase of each cell (3.14±1.17 rad, indicated by the dashed red line; circular median test, p<0.001); **[G]** Unwrapped theta phase by spike number within theta spike trains, averaged across all cells (error bars indicate circular standard deviation)

In accordance with previous studies, and similar to the changes in ripple band firing phase within each candidate replay event described above, the theta phase shift between successive spikes was consistently negative (Figure 2C). Specifically, the distribution of circular mean phase shifts across theta n-spike trains was non-uniform, with an overall circular mean that differed significantly from zero. Importantly, this effect was true when each spike pair (i.e. first, second, third etc.) was considered separately (Figure 2D). Similarly, the mean slope of the relationship between normalised time within the n-spike train and theta phase was consistently negative and significantly different from zero across cells (Figure 2E). Finally, the distribution of intercepts was also non-uniform, with a circular mean value that was significantly later than the phase of peak firing in the theta cycle (Figure 2F). These results indicate that the theta band phase of putative place cell firing during active movement begins after the phase of peak firing in the theta cycle and becomes progressively earlier as the spike train continues (Figure 2G). Conversely, consistent changes in theta band firing phase were not observed across putative principal neurons recorded simultaneously in the deeper layers of MEC, nor if we restricted that analysis to just grid cells (see Figure S7 for details).

Next, we directly compared within-event time vs. phase relationships between candidate replay events during REST and theta n-spike trains during RUN across cells. Mean firing rates were highly correlated between REST and RUN (r=0.507, p<0.001), although firing rates were significantly higher during RUN (0.458±0.728 vs. 0.209±0.6.0Hz, t(959)=11.9, p<0.001). Across cells that participated in n-spike events in both sessions (n=853), phase shifts per spike pair were greater in the ripple band during REST than the theta band during RUN (−0.941±1.18 vs -0.309±0.582 rad; Watson-Williams test, p<0.001), but there were fewer spikes per event (median±SD=3±0.467 vs 4±1.45; t(852)=-24.2, p<0.001), such that normalised time vs. phase slopes (equivalent to the total change in firing phase across each spike train) were less negative overall (−0.502±2.66 vs -1.44±2.77 rad; t(852)=6.18, p<0.001). Interestingly, within-event time vs. phase slopes were correlated across cells between sessions (r=0.0713, p<0.05), indicating that cells which exhibited larger changes in ripple band firing phase across n-spike trains during individual replay events tended to exhibit larger changes in theta firing phase across individual n-spike trains during active movement on the track.

Crucially, previous studies have demonstrated that place cell theta firing phase during movement does not simply change as a function of spike number or time since firing began but is most strongly modulated by distance travelled through the place field (O’Keefe and Recce, 1993). As a result, theta firing phase encodes information about the animal’s location, beyond that provided by firing rates alone (Jensen and Lisman, 2000; Huxter et al., 2003). In addition, theta firing phase encodes information about movement direction, even when this conflicts with the head direction signal (Cei et al., 2014; Maurer et al., 2014; Raudies et al., 2015; Bush and Burgess, 2020). Next, we sought to characterise this aspect of theta phase precession in our data, and then establish whether ripple band firing phase shared a similar relationship with decoded location within the place field across replay events.

First, we defined place fields as ≥10 contiguous 2cm spatial bins with smoothed firing rate greater than the mean across all bins and a peak firing rate of ≥1Hz, with 919/1044 putative pyramidal cells (88%) active during RUN having ≥1 field on either outbound or inbound runs along the track. Next, we identified a total of 1355/1499 place fields on outbound and 1256/1423 place fields on inbound runs along the track that passed our criteria for inclusion in phase precession analyses (≥5 spikes fired within the field on all runs through that field, covering ≥50% of the place field), with 873/1044 (83.6%) cells having ≥1 field included on either outbound or inbound runs along the track (see Figure S8, Methods for details). Finally, we characterised the relationship between normalised distance travelled through the field (collapsed across all runs through the field in each session) and theta firing phase using circular-linear regression. Again, given that place cells tended to fire >1 spike during each oscillatory cycle, unlike during replay, we restricted these analyses to the first spike from each theta cycle.

As expected, the median slope of this relationship was consistently negative and significantly different from zero across cells (Figure 3A), with no difference in slopes between the outbound and inbound maps (across cells that had ≥1 field during runs in each direction, t(537)=1.43, p=0.15; outbound=-2.78±5.24, inbound=-3.07±4.89 rad/field). In addition, the distribution of normalised location vs. theta firing phase intercepts was non-uniform, with a circular median value that was significantly later in the theta cycle than the phase of peak firing (Figure 3B). As such, place cell firing typically shifted from late to early theta phases as the firing field was traversed (Figure 3C). Conversely, no such relationship was observed across putative excitatory cells in the deeper layers of MEC, nor in grid cells specifically (see Figure S8 for details).

**Figure 3:**
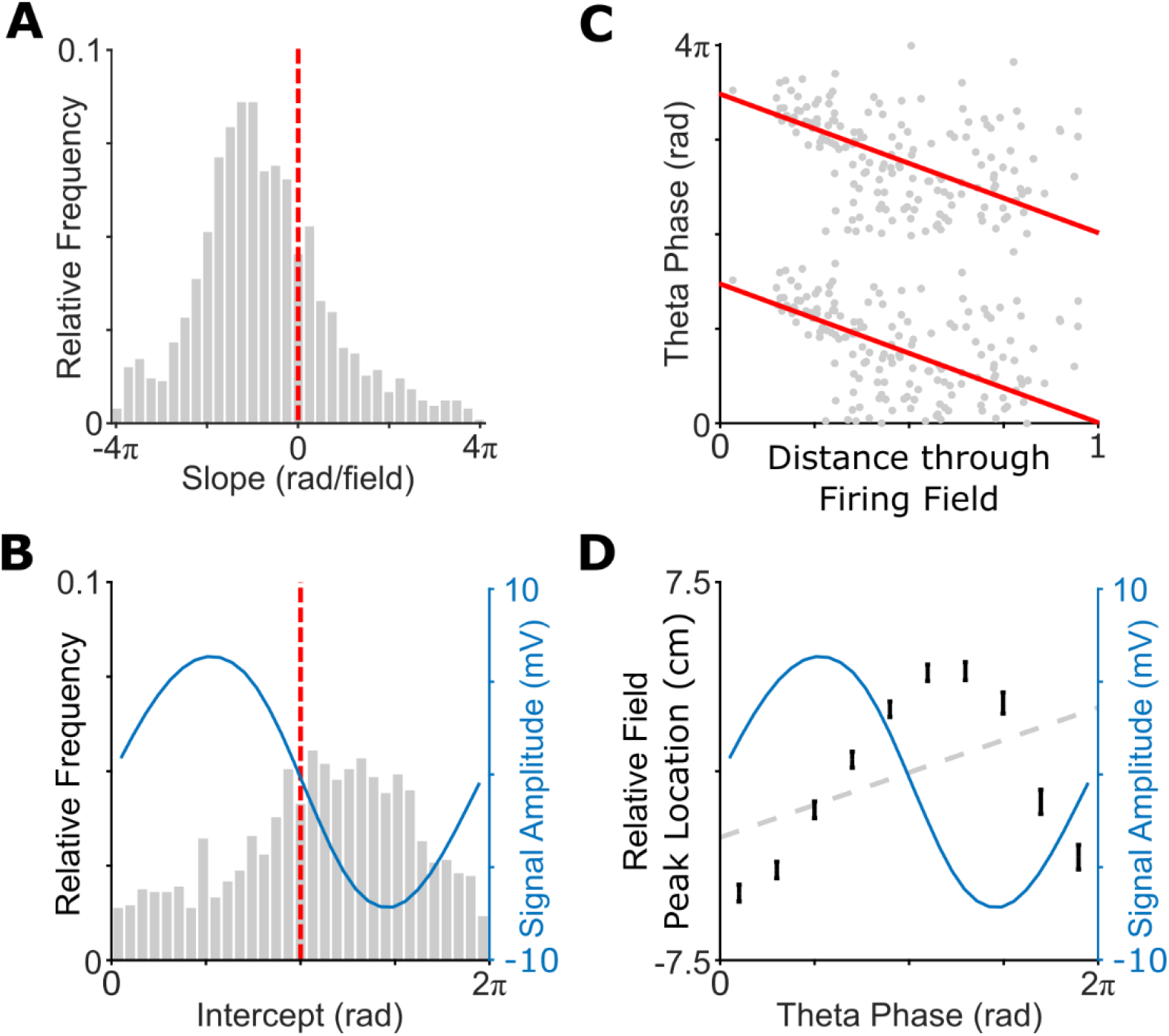
Relationship between actual location and theta firing phase during movement. **[A]** Distribution of mean normalised location vs. theta phase slopes (overall median±SD=-2.77±3.94 rad/field), which differs significantly from zero (t(856)=-18.0, p<0.001); **[B]** Distribution of mean normalised location vs. theta phase intercepts across cells (overall circular median±circular SD=3.90±1.63 rad). This distribution is non-uniform (Rayleigh test, z=99.6, p<0.001) with a median value that differs significantly from the phase of peak firing (3.20±1.17 rad, indicated by a dashed red line; circular median test, p<0.001); **[C]** Relationship between distance travelled through the firing field and theta phase for a typical place field, alongside the circular-linear fit (red line); **[D]** Overall theta sweep, showing median relative distance to the peak of the nearest firing field (black error bars) across all active place cells, alongside actual location (dashed grey line) and mean signal amplitude (blue line) in each of 10 theta phase bins

Coordinated phase precession across place cells generates ‘theta sweeps’ of activity during movement, with place cells encoding for locations behind the animal being active early, and place cells encoding for locations ahead of the animal being active later, in each theta cycle (Burgess et al., 1994; Skaggs et al., 1996; Johnson and Redish, 2007; Feng et al., 2015; Kay et al., 2020; Wang et al., 2020). To examine this phenomenon in our data, we split each theta cycle into ten discrete phase bins and computed the relative distance between the animal’s average location in that cycle and the peak of the closest place field for all cells that were active in each phase bin (again, including only the first spike fired by each cell in each cycle). To facilitate subsequent comparison with replay events, we then averaged those data across all theta cycles in each continuous period of movement (defined as running speed ≥10cm/s for a duration of ≥1s) and used linear regression to estimate the distance covered by that average theta sweep (see Methods). This revealed an average forward sweep covering 13.2±51.4cm within each theta cycle (Figure 3D), which was significantly different from zero across events (t(5441)=17.3, p<0.001). These forward sweeps exhibited an equivalent movement speed of 108±504cm/s, which significantly exceeds the animal’s actual movement speed of 43.6±16.7cm/s during the same events (t(5441)=10.9, p<0.001). Crucially, theta sweeps also had a positive slope (i.e. consistent with movement direction) in 68.5% of all events (more than expected by chance, binomial test, p<0.001), indicating that they reliably encoded the animal’s current trajectory.

To establish if a similar relationship existed between decoded location within a place field during replay and ripple band firing phase, we next sought to identify which candidate replay events encoded coherent spatial trajectories on the Z-maze. To do so, we used a Bayesian decoding algorithm and trajectory fitting procedure (Olafsdottir et al., 2015; see Methods for further details). This revealed that 13.9% of all candidate events (3523 / 25328, 13.5±4.7% across sessions, range 4.21-24.9%) corresponded to the replay of linear trajectories along the track, with events that were longer, incorporated more active cells or more spikes being more likely to show significant linear replay (see Figure S9 for details). Interestingly, there was no difference in the magnitude of within-event ripple band phase shifts per spike pair (circular median test, p=0.54; significant=-0.936±1.50, non-significant=-0.882±1.13 rad/spk) or within-event time vs. ripple band firing phase slopes (t(779)=-0.048, p=0.96; significant-0.559±4.30, non-significant=-0.506±2.37 rad) between significant and non-significant linear replay events across cells.

Next, we used these fitted trajectories to estimate the mean decoded location in each ripple band cycle during significant linear replay events. For each place field, we could then examine the relationship between ripple band firing phase across all significant decoded trajectories that passed through that firing field in the same direction of movement (i.e. considering forward and reverse replay events separately), and decoded location within the firing field in the corresponding ripple band cycle. First, we identified a total of 1569 place fields that passed our criteria for inclusion (≥5 spikes fired within the field across all significant decoded trajectories through that field, covering ≥50% of the place field), with 576/1025 cells (56.2%) having ≥1 field included on either the outbound or inbound rate maps (see Methods for details). Second, we characterised the relationship between normalised location within the field, according to the decoded trajectory, and ripple band firing phase using circular-linear regression.

Remarkably, we found that the mean slope of this relationship was consistently negative and significantly different from zero across cells (Figure 4A). There was no difference in slopes between fields on the outbound and inbound maps, for cells with fields included on both maps (t(277)=-0.361, p=0.718; outbound=-0.201±6.09, inbound=-0.437±6.03 rad/field); or by forward or reverse trajectories through these fields, for cells with sufficient spikes fired in each decoded movement direction through the same field (t(343)=-0.825, p=0.41; forward=-0.556±6.03, reverse=-0.201±6.07 rad/field). In addition, the distribution of decoded location versus ripple band firing phase intercepts was non-uniformly distributed, with a circular median value that was significantly later in the ripple cycle than the mean firing phase of all spikes fired (Figure 4B). As such, place cell firing typically shifted from late to early ripple band phases as replay trajectories passed through the firing field (Figure 4C).

**Figure 4:**
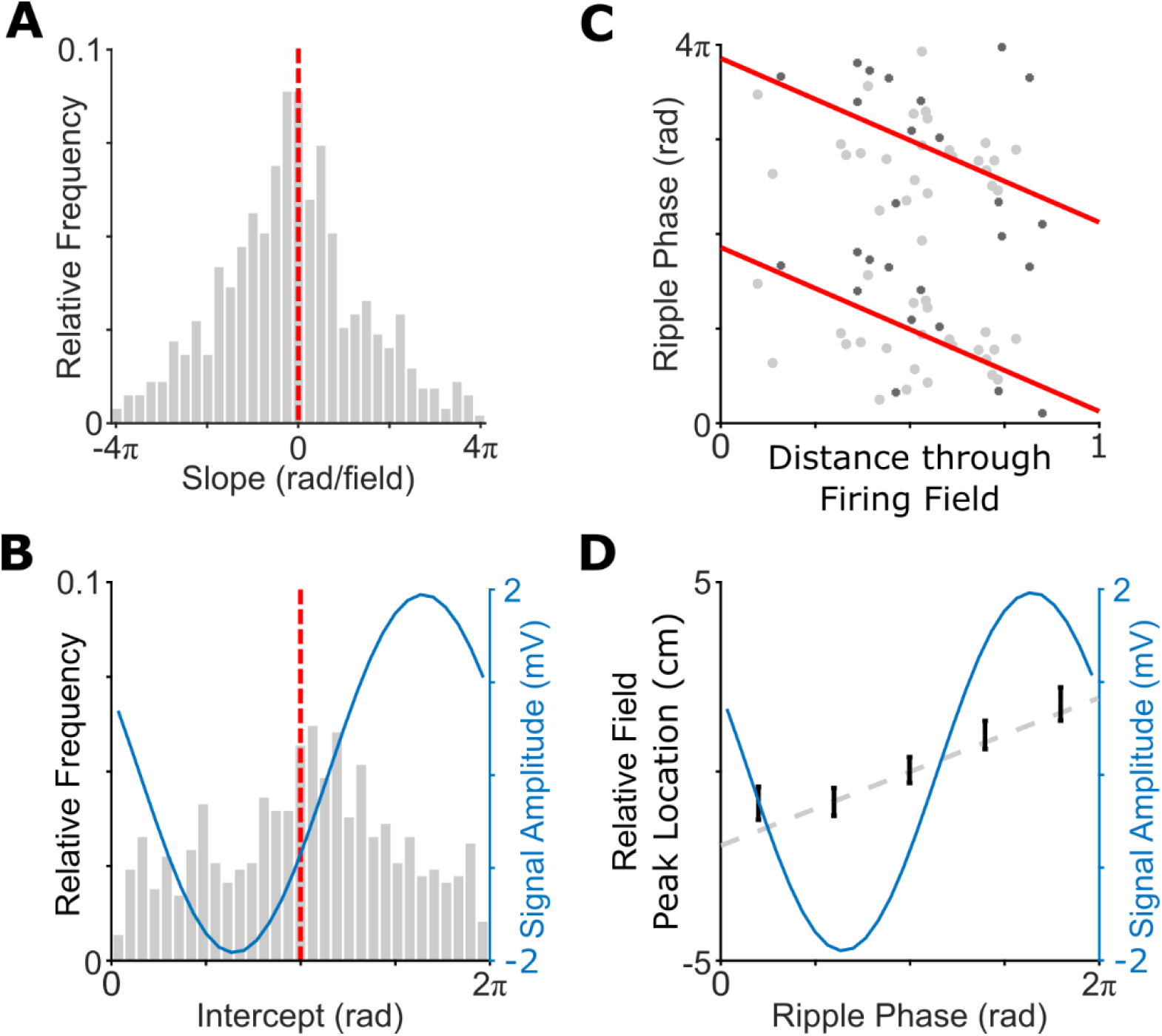
Relationship between decoded location and ripple band firing phase during replay. **[A]** Distribution of mean normalised location vs. ripple band phase slopes (overall median±SD=-0.505±4.80 rad/field), which differs significantly from zero (t(553)=-2.36, p=0.0184); **[B]** Distribution of mean normalised location vs. ripple band phase intercepts across cells (overall circular median±circular SD=3.42±1.78 rad). This distribution is non-uniform (Rayleigh test, z=24.0, p<0.001) with a median value that differs significantly from the phase of peak firing (3.12±0.61, indicated with a dashed red line; circular median test, p<0.001); **[C]** Relationship between decoded location within the firing field and ripple band firing phase for the place field shown in Figure 3C, incorporating spikes from both n-spike trains (light grey) and non-n-spike trains (dark grey), alongside the circular-linear fit (red line); **[D]** Overall ripple sweep, showing median relative distance to the peak of the nearest firing field across all active place cells alongside decoded location (dashed grey line) and mean signal amplitude (blue line) in each of 5 ripple band phase bins

Importantly, there was no difference in the slope of decoded location versus ripple band firing phase between events in which each place cell fired only a single spike and those in which each place cell fired ≥3 spikes, across cells with sufficient in-field spikes in both cases (t(150)=0.158, p=0.875; single spike events=-0.478±6.90, n-spike events=-0.614±5.77 rad/field; see Figure 4C for an example). This indicates that, even when a place cell fired only a single spike within its firing field during a significant replay trajectory, it did so at a ripple band phase that indicated location within that firing field. Moreover, when analysed separately, a negative relationship between decoded location within the place field and ripple band firing phase was observed across cells in all but one animal (range of - 1.31 to 0.16 rad/field, t(5)=-2.99, p<0.05). Quantitatively similar results were observed during online replay events, which occurred while the animal was awake but immobile at the corners of the track, suggesting that this phenomenon is preserved across behavioural states (Figure S10A). However, no such relationship was observed across putative excitatory cells in the deeper layers of MEC, nor across grid cells in that region specifically (Figure S10B, C).

Next, we asked if coordinated ripple band phase precession across place cells during offline replay events also gave rise to ripple band sweeps of activity within each oscillatory cycle, analogous to the theta sweeps observed during active movement. To address this, we split each ripple band cycle into five discrete phase bins and computed the relative distance between the animal’s average decoded location in that cycle and the peak of the closest place field for all cells that were active in each phase bin. We then averaged these data across all ripple band cycles in each significant replay event and used linear regression to estimate the distance covered by that average ripple band sweep (see Methods). This revealed an average forward sweep covering 2.61±56.3cm within each ripple cycle (Figure 4D), which was significantly different to zero across events (t(3404)=4.29, p<0.001). In addition, the sequence of locations encoded by place cell firing across phase bins was consistent with the movement direction of the decoded replay trajectory in 54.1% of all events, indicating that ripple band phase coding could be used to infer movement direction more often than expected by chance (binomial test, p<0.001; Zutshi et al., 2017; Bush and Burgess, 2020). Interestingly, however, these forward sweeps exhibited an equivalent movement speed of 6.02±134m/s, which is not significantly different than decoded movement speed during the same events (t(3404)=-0.532, p=0.594). Unlike theta sweeps, therefore, ripple band sweeps do not appear to have a strong prospective component.

Finally, we directly compared within-field location vs. phase relationships between theta and ripple band oscillations during RUN and REST, respectively, across n=561 cells with ≥1 place field that passed our criteria for inclusion in each session. During RUN, firing fields incorporated significantly more spikes (67.0±51.7 vs 14.7±15.4; t(560)=-29.5, p<0.001) and median phase precession slopes were significantly more negative (−2.77±3.94 vs -0.505±4.80 rad/field; t(560)=8.0, p<0.001). In this case (unlike the correlation between within-event time vs. phase relationships for n-spike cells), we found no correlation between within-field location vs. phase slopes aggregated over multiple trajectories through each place field during RUN and REST (r=-0.0399, p=0.346). We also found no evidence for any correlation in within-field location vs. phase slopes between firing fields of the same cell on outbound and inbound runs during either REST or RUN, or between ripple band phase precession slopes between forward and reverse trajectories through the same firing fields during REST (all p>0.19). This suggests that there is no relationship between within-field location vs. phase slopes across cells, either between behavioural states, between firing fields of the same cell, or between different movement directions during replay.

## Discussion

We have demonstrated that the phase code for location exhibited by place cells during movement related theta oscillations is preserved during ripple band activity, when the hippocampus is believed to replay information to the neocortex. Specifically, we have shown that place cells which fire multiple spikes during candidate replay events do so at progressively earlier phases of the ongoing ripple band oscillation; that there is a consistently negative relationship between decoded location within the place field and ripple band firing phase across all replay trajectories that pass through the field; and that this is associated with forward sweeps of activity within each ripple cycle, which could be used to infer movement direction at the population level. Importantly, these results are consistent across online and offline replay events (i.e. those recorded during quiescent waking periods on the track and during subsequent rest, respectively), indicating that they are not dependent on behavioural state. In addition, the relationship between decoded within-field location and ripple band phase is consistent across events where cells fire only one spike or multiple spikes, indicating that it does not simply arise from changes in ripple band firing phase across multiple spikes within single events. In sum, this demonstrates that hippocampal phase coding is not restricted to place cell firing during active movement; nor to sustained low frequency oscillations with relatively constant rhythmicity, consistent with a growing body of experimental work across species (Jaocbs, 2014; Zheng et al,. 2016; Eliav et al., 2018; Bush and Burgess, 2020; Qasim et al., 2020).

These findings have two major implications for our understanding of neural coding and the function of the mammalian hippocampus. First, they suggest shared or similar mechanisms for updating place cell activity according to real and simulated movement trajectories during theta and ripple band oscillations, respectively. Specifically, the phase coding mechanism hypothesised to update place cell activity during replay (Hasselmo, 2009; Bicanki and Burgess, 2018; Yu et al., 2020) might be the same as that used to update place cell activity according to path integration during actual movement (Mehta et al., 2002; Burgess et al., 2007). Consistent with a relationship between theta and ripple band phase precession, we found that the slope of within-event time vs. phase relationships during individual candidate replay events and theta n-spike trains were correlated across cells. Interestingly, however, the LFP phase of peak firing differs by ∼π rad between theta and ripple band oscillations, potentially suggesting a different spatial distribution of LFP sources with a common mode of phase coding. Second, the phase coding of external and / or internal variables may reflect a more general mechanism for transmitting information from the hippocampus to downstream circuits. The theta phase code for distance travelled within the firing field improves the accuracy of location decoding (Jensen and Lisman, 2000) and allows movement direction to be inferred from population activity in each oscillatory cycle, which is not always possible using head direction cells (Cei et al., 2014; Maurer et al., 2014; Raudies et al., 2015; Zutshi et al., 2017; Bush and Burgess 2020). More generally, the multiplexing of information in firing rate and phase could allow for much richer coding of task relevant variables in the human brain across a range of network states and cognitive domains (Kurth-Nelson et al., 2015; Heusser et al., 2016; Liu et al., 2020; Qasim et al., 2020; Vaz et al., 2020).

Interestingly, we found no evidence for consistent changes in theta or ripple band firing phase in putative excitatory neurons recorded simultaneously in the deeper layers of MEC, nor across a sub-population of grid cells specifically. This suggests that ripple band phase precession may only be observed in cells that also exhibit theta phase precession. Future work might therefore look for ripple band phase precession in grid cells from the superficial layers of MEC, which are known to display robust theta phase precession and engage in replay events (Hafting et al., 2008; Olafsdottir et al., 2016; O’Neill et al., 2018). It has yet to be established whether large populations of grid cells in the deeper layers of MEC exhibit theta phase precession on linear tracks (Hafting et al., 2008), although previous studies have described this phenomenon in open field recordings (Climer et al., 2013; Jeewajee et al., 2013), in contrast to our results on the Z-shaped track. Elsewhere, recent studies have demonstrated a link between theta sweeps during active exploration and enhanced replay of trajectories through that environment (Feng et al., 2015, Drieu et al. 2018; Farooq et al., 2019; Muessig et al., 2019). Hence, future work should examine the development with experience of coordinated theta phase precession across place cells during active exploration and ripple band phase precession during subsequent replay trajectories. Given that we found equivalent ripple band phase precession during forward and reverse replay events, despite the fact that place cell firing has never occurred during reverse movement through the corresponding place field, this work could specifically investigate the relationship between the retrospective component of theta sweeps during active movement and subsequent ripple band phase precession during reverse replay events (Wang et al., 2020).

One important question raised by these findings is whether downstream targets of the hippocampus could feasibly recover information encoded in ripple band firing phase. The duration of ripple band oscillatory cycles is on the order of ∼5ms, and the phase range of ripple band precession is significantly less than that observed during theta oscillations, on the order of milliseconds and similar to the spike width of a typical pyramidal cell. It is not clear if downstream neurons can distinguish inputs on a timescale of milliseconds, although we note that this is close to the timescale of integration in a typical cortical neuron; that coincidence detection may be facilitated by the large number of hippocampal neurons that are active in each candidate replay event (∼25% on average); and that cortical dendrites are capable of distinguishing temporal input patterns on a similar timescale (Branco et al., 2010). This question could be addressed, in future, by computational modelling, or by identifying methods for selectively disrupting the ripple band phase code and observing the effects on behaviour (Robbe and Buzsaki, 2009). In addition, future work should explore the differences between theta and ripple band forward sweeps – specifically, that the former consistently exhibit a strong prospective component, while the latter do not. The encoded direction and speed of ripple band sweeps described here is highly variable, and while some of this variability is undoubtedly due to sampling error, it is also possible that the properties of ripple band sweeps are modulated by behavioural variables (Olafsdottir et al., 2017).

In summary, we have demonstrated that the phase code for location exhibited by place cells is not specific to active movement or the theta frequency band. Consistent with a growing body of research, these findings suggest that the hippocampus might employ similar mechanisms to generate and project phase coded information to downstream targets across a range of behavioural states.

## Acknowledgements

The authors wish to thank Robin Hayman for his input during the preparation of this manuscript. FO is supported by a Donders Mohrmann fellowship, and this research was also funded in part by the Wellcome Trust [Grant number 202805/Z/16/Z to NB and 212281/Z/18/Z to CB]

## Author Contributions

Project administration, DB; Supervision, NB and CB; Writing – original draft preparation, DB; Writing – review and editing, DB, HFO, CB and NB; Investigation, HFO and CB; Methodology, DB, HFO, CB and NB; Funding acquisition, HFO, CB and NB

## Methods

### Surgery

Six male Lister Hooded rats (330–400 g at implantation) received two microdrives, each carrying eight tetrodes of twisted 17μm HM-L coated platinum iridium wire (90% and 10%, respectively; California Fine Wire), targeted to the right CA1 (ML: 2.2 mm, AP: 3.8 mm posterior to bregma) and left medial entorhinal cortex (MEC; ML=4.5 mm, AP=0.3–0.7 anterior to the transverse sinus, angled between 8–10°). Wires were platinum plated to reduce impedance to 200–300 k at 1 kHz. After rats had recovered from surgery they were maintained at 90% of free-feeding weight with ad libitum access to water and were housed individually on a 12-h light-dark cycle. All procedures were approved by the UK Home Office, subject to the restrictions and provisions contained in the Animals (Scientific Procedures) Act of 1986.

Screening was performed post-surgically after a 1-week recovery period. An Axona recording system (Axona Ltd.) was used to acquire single units and position data (for details of the recording system and basic recording protocol see Barry et al., 2007). Position and head direction were inferred using an overhead video camera to record the location of two light-emitting diodes (LED) mounted on the animals’ head-stages at a sample rate of 50 Hz. Tetrodes were gradually advanced in 62.5μm steps across days until place cells (CA1) or grid cells (MEC) were found. EEG data were concurrently acquired at a sample rate of 4800Hz.

### Behavioural Protocol

The experiment was run during the animals’ light period, to facilitate rest during the REST session. During RUN sessions animals shuttled back and forward on a Z-shaped track comprised of 10cm wide runways covered with black paint, raised 75cm off the ground. The two parallel sections of the Z (190cm each) were connected by a diagonal section (220cm; see Figure 1A). The entire track was surrounded by plain black curtains. Animals were pre-trained to run on the track, taking between 3 and 6d before they would shuttle fluently from one end to the other. At the start of each session, rats were placed at one end of the Z-track. The same end was used as a starting location for every day of the experiment and for every rat. The ends and corners of the track were baited with sweetened rice to encourage running from one end to the other.

Following the RUN session, rats were placed in the REST enclosure for an hour and a half. The rest enclosure consisted of a cylindrically shaped environment (18cm diameter, 61cm high) with a towel placed at the bottom and was located outside of the curtains that surrounded the Z-track. Animals were not able to see the surrounding room while in the rest enclosure. Prior to recording, rats had been familiarized with the rest environment for at least 7d. Following the REST session, rats foraged in a familiar open field environment for 20 min, to allow functional classification of MEC cells (Olafsdottir et al., 2016).

### Identifying Putative Principal Cells, Grid Cells

All analyses were restricted to putative principal cells, identified by manual inspection of waveforms and mean firing rates ≤10Hz across the entire recording session. We classified MEC cells as grid cells using a shuffling procedure similar to that applied elsewhere (Wills et al., 2010). Specifically, we generated firing rate maps for each cell during the 20 minute foraging session that followed REST. We then computed both the standard (Sargolini et al., 2006) and modified (Langston et al., 2010) gridness measures for each cell and compared those values with a null distribution generated by randomly permuting the spike train of each cell relative to the tracking data by a temporal distance of ≥30s, 100 times. Grid cells were subsequently classified as those for which the standard or modified gridness scores exceeded the 97.5th percentile of the matching null distribution. This identified 80/1112 grid cells during RUN and 76/877 during REST.

### Identifying Candidate Replay Events

We used two methods to identify candidate replay events – either as periods of elevated multi-unit activity (MUA) or ripple band power. In the former case, we first generated a MUA histogram for all putative principal cells in 1ms time bins. This histogram was then smoothed using a Gaussian kernel with 5ms standard deviation and candidate replay events identified as periods of Z≥0 with peak Z≥3. Any candidate events separated by ≤40ms were merged and any remaining events with a duration of ≤40ms discarded. Finally, we excluded any events during which <5 or 15% of all pyramidal cells were active (whichever is the larger), median running speed >10cm/s, or duration >0.5s.

In the latter case, we first identified the EEG channel in each RUN session with the highest 6-12Hz theta signal to noise ratio. Next, we filtered LFP data from that channel in the 150-250Hz ripple band using a 400^th^ order finite impulse response (FIR) filter, extracted the amplitude of the filtered signal using the Hilbert transform and smoothed that time series using a Gaussian kernel with 5ms standard deviation. Candidate replay events were then identified as periods with ripple band amplitude Z≥0 and peak ripple band amplitude Z≥3. Any candidate events separated by ≤40ms were merged and any remaining events with a duration of ≤40ms discarded. Finally, we excluded any events during which <5 or 15% of all pyramidal cells were active (whichever is the larger), median running speed >10cm/s, or duration >0.5s.

### Changes in Ripple Band Firing Phase within Candidate Replay Events

To examine changes in the ripple band firing phase of active cells over the course of candidate replay events, we first identified the EEG channel in each RUN session with the highest 6-12Hz theta signal to noise ratio. Next, we filtered LFP data from that channel in the 150-250Hz ripple band using a 400^th^ order FIR filter, extracted the phase of the filtered signal at each time point using the Hilbert transform, and computed the preferred ripple band firing phase of each cell as the circular mean firing phase across all candidate replay events. We then adjusted LFP and firing phase values such that the circular mean preferred firing phase across all putative principal cells in each session was equal to π rad.

In addition, we quantified the ripple band phase modulation of firing for each cell as the resultant vector length of ripple band firing phases across all candidate replay events (referred to as the ‘phase locking value’). To establish the significance of phase modulation, we compared the true resultant vector length with a surrogate distribution generated by shuffling spike times for that cell relative to LFP ripple band phase by a temporal distance of ≥30s, 1000 times, and setting the threshold for significance as the 99^th^ percentile of that distribution. For illustration purposes, we also generated circular firing rate histograms for each cell using 30 equally sized phase bins.

Next, we identified all cells in each candidate event that fired ≥3 spikes and computed both the circular mean phase shift between successive spikes; and the circular-linear correlation between normalised spike time (i.e. where the first spike fired by that cell in that replay event occurs at *t*=0 and the last at *t*=1) and ripple band firing phase (Figure 1B), for each candidate event (Kempter et al., 2012). The latter provides an intercept and slope of the circular-linear relationship between normalised spike time and ripple band phase relationship for that cell in that event. We subsequently computed the average phase shift, slope and circular mean intercept across all events for each cell. In addition, we repeated the analysis above but considered only the first spike fired in each oscillatory cycle, in which case only cells that fire in ≥3 cycles were included.

### Changes in Theta Firing Phase within N-spike Trains

To examine changes in the theta firing phase of active cells over the course of n-spike trains, we first generated a spike train histogram for each cell with a bin size of 5ms using only spikes fired at a running speed of ≥10cm/s. This histogram was smoothed using a Gaussian kernel with 80ms standard deviation, converted to a firing rate, and candidate theta spike trains identified as contiguous periods with a mean firing rate ≥1Hz. Any candidate events separated by ≤500ms were merged and any remaining events with duration <500ms discarded. Finally, we excluded any events that incorporate <3 spikes.

Next, we filtered LFP data in the 6-12Hz theta band using a 400^th^ order FIR filter, extracted the phase of the filtered signal at each time point using the Hilbert transform, and computed the preferred theta firing phase of each cell as the circular mean firing phase across all candidate theta spike trains. We then adjusted LFP and firing phase values such that the circular mean preferred firing phase across all putative principal cells in each session was equal to π rad.

In addition, we quantified the theta phase modulation of firing for each cell as the resultant vector length of theta firing phases across all spikes fired at a running speed of ≥10cm/s (referred to as the ‘phase locking value’). To establish the significance of phase modulation, we compared the true resultant vector length with a surrogate distribution generated by shuffling spike times for that cell relative to LFP theta phase by a temporal distance of ≥30s, 1000 times, and setting the threshold for significance as the 99^th^ percentile of that distribution. For illustration purposes, we also generated circular firing rate histograms for each cell using 30 equally sized phase bins.

Finally, we computed both the circular mean phase difference between successive spikes in each theta spike train; and the circular-linear correlation between normalised spike time (i.e. where the first spike fired by that cell in that theta spike train occurs at *t*=0 and the last at *t*=1) and theta firing phase (Figure 2A). This provided an intercept and slope for the circular-linear relationship between normalised spike time and theta phase for that cell in that event. We subsequently computed the average phase shift, slope and circular mean intercept across all events for each cell. In addition, we repeated the analysis above but consider only the first spike fired in each theta cycle, in which case only cells that fire in ≥3 cycles are included.

### Theta Phase Precession

To assess theta phase precession during running on the track, we first generated firing rate maps for each active cell in each session. To do so, we excluded spikes fired in areas where the animals regularly performed non-perambulatory behaviours (for example, eating or grooming) - specifically, in the final 10 cm at either end of the track and 5 cm around each of the two corners – and from ‘unsuccessful’ runs (i.e. those in which the animal did not progress from one end of the Z track, along all three sections to the other end). We then linearised each animal’s path and computed dwell time and the total number of spikes fired at running speeds of ≥10cm/s in 2cm bins. These histograms were each smoothed using a Gaussian kernel with standard deviation of 5 bins, and firing rates computed by dividing the spike number by dwell time. We generated separate rate maps for runs in the outbound and inbound directions, and defined place fields for each cell as contiguous regions of at least 10 spatial bins with firing rate Z≥0 and a peak in-field firing rate of ≥1Hz.

Next, we filtered LFP data in the 6-12Hz theta band using a 400^th^ order FIR filter and extracted the phase of the filtered signal at each time point using the Hilbert transform. We then extracted the theta firing phase of all spikes fired by each cell within each firing field at running speeds of ≥10cm/s, and computed the preferred firing phase of each cell as their circular mean firing phase. We then adjusted LFP and firing phase values such that the circular mean preferred firing phase across all putative principal cells in each session was equal to π rad.

Next, we computed the circular-linear correlation between the relative distance travelled through the field and theta firing phase for any field with ≥5 spikes that covered ≥50% of the place field. Importantly, we included only the first spike fired in each theta cycle in this analysis. This provided an intercept and slope of the circular-linear relationship between actual location within the firing field and theta firing phase relationship for each place field. We subsequently computed the average slope and circular mean intercept across all fields for each cell.

### Decoding Location on the Track

Before decoding location during candidate replay events, we first used a Bayesian decoding framework to establish the accuracy with which location on the track could be decoded from activity across a population *N* of place cells during each RUN session (Zhang et al., 1998). To do so, we computed the number of spikes *k* fired by each cell *i* in consecutive *T*=500ms time bins during successful outbound and inbound runs when the running speed of the animal was ≥10cm/s. Using the firing rate maps *α*_*i*_(x) for each cell on outbound and inbound runs, respectively, we then computed a posterior probability distribution *P*(*x* | *K*)describing the likelihood that the animal was located in each 2cm bin of the firing rate map, given the population spiking activity, according to Equation 1. The true location in each time bin was taken as the average location across all samples within that bin, and the decoded location was taken as the spatial bin with the greatest probability across all spatial bins. This allowed a decoding error to be computed for each time bin, and a median error across all time bins to be computed for outbound (13.7cm) and inbound (12.8cm) runs in each session, respectively. As expected, the mean decoding error for each session (averaged across runs in each direction) was strongly negatively correlated with the total number of pyramidal cells recorded (r=-0.65, p<0.001).

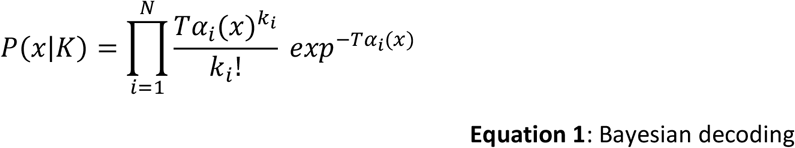

### Decoding Location during Replay Events

Next, we used similar methods to decode location from place cell population activity during candidate replay events. In that case, however, we computed the number of spikes *k* fired by each cell *i* in consecutive time bins of *T*=10ms throughout the replay event, starting at the time of the first spike fired by any cell. We then decoded location independently in each time bin where at least one spike was fired by any cell using the Bayesian framework described by Equation 1 (note that this includes all putative pyramidal cells, whether or not they have a place field on the track in either running direction). This provides a spatial bin x time bin posterior probability matrix for each candidate event, with probability values in each time bin normalised so that they sum to one.

We then identified the linear fit thorough that posterior probability matrix which maximises the summed probability under a line of 15 spatial bins in width (i.e. 30cm), testing all potential spatial bins as a starting point and movement speeds of 100 : 50 : 5000 cm/s in either forward or reverse directions along the track (Davidson et al., 2009). We recorded the intercept (i.e. spatial origin of the decoded trajectory) and slope (i.e. running speed of the decoded trajectory) of that line, as well as the average summed probability per time bin (or ‘fit score’). To establish the significance of each decoded trajectory, we shuffle the identity of active cells within each event 100 times (noting that our minimum threshold of 5 active cells allows a minimum of 120 unique shuffles) and re-computed the optimal fit score for each shuffle. An event is considered to be significant if the true fit score exceeds the 95^th^ percentile of the fit score distribution obtained by shuffling. As expected, there is a strong positive correlation between the total number of pyramidal cells recorded in a session and the overall proportion of significant events (r=0.646, p<0.001).

### Ripple Band Phase Precession

To examine the relationship between decoded location within a firing field and ripple band firing phase, we first use the linear trajectories identified above to compute the average decoded location in each ripple band cycle in each significant replay event. We use these decoded locations to identify all spikes fired by each cell within each firing field across all significant replay events in each session. We then compute the circular-linear correlation between the relative distance of each spike through the field (given the decoded movement direction) and ripple band firing phase for any field with ≥5 spikes that cover ≥50% of the place field. This provided an intercept and slope for the circular-linear relationship between decoded location within the firing field and ripple band firing phase for each place field. We subsequently computed the average slope and circular mean intercept across all fields for each cell.

### Theta and Ripple Band Sweeps

To examine theta sweeps during movement on the track, and facilitate comparison with individual replay events, we first identify ‘movement events’: periods of continuous movement with running speed ≥10cm/s and minimum duration ≥1s. For each active cell in each movement event, we extract the theta phase at which the first spike was fired during each oscillatory cycle, and the relative distance between the average true location of the animal in that theta cycle and the peak of the nearest place field of that cell. Next, we compute the median relative distance across all spikes fired by all cells in all cycles within that event for ten equally sized theta phase bins, to provide an estimate of the average theta sweep for that event. Finally, we used linear regression to estimate the slope of the relationship between theta phase and decoded location, restricting our analyses to theta phase bins that showed forward movement in the grand average plot (Figure 3C, specifically, between 0 and 7π/5 rad). These slopes were subsequently used to estimate the overall range and corresponding speed of theta sweeps for each event.

To examine ripple band sweeps during sleep, we extract the ripple band firing phase of all spikes fired by all active cells in each significant replay event, and compute the relative distance between the average decoded location in that ripple band cycle (estimated using the slope and intercept of fitted trajectories) and the peak of the nearest place field of that cell. Next, we compute the median relative distance across all spikes fired by all cells in all cycles within that event, for five equally sized ripple band phase bins, to provide an estimate of the average ripple sweep for that event. Finally, we used linear regression to estimate the slope of the relationship between ripple band phase and decoded location, restricting our analyses to ripple band phase bins that showed forward movement in the grand average plot (Figure 4C, specifically, between 2π/5 and 2π rad). These slopes could subsequently be used to estimate the overall range and corresponding speed of theta sweeps for each event.

## Code Availability

All analysis was carried out using Matlab (Mathworks, Natick MA) and all code is available from the authors upon reasonable request. All circular statistics were computed using CircStat: A MATLAB Toolbox for Circular Statistics (Berens, 2009).

