## Supplementary Figures for "Ripple Band Phase Precession of Place Cell Firing during Replay"

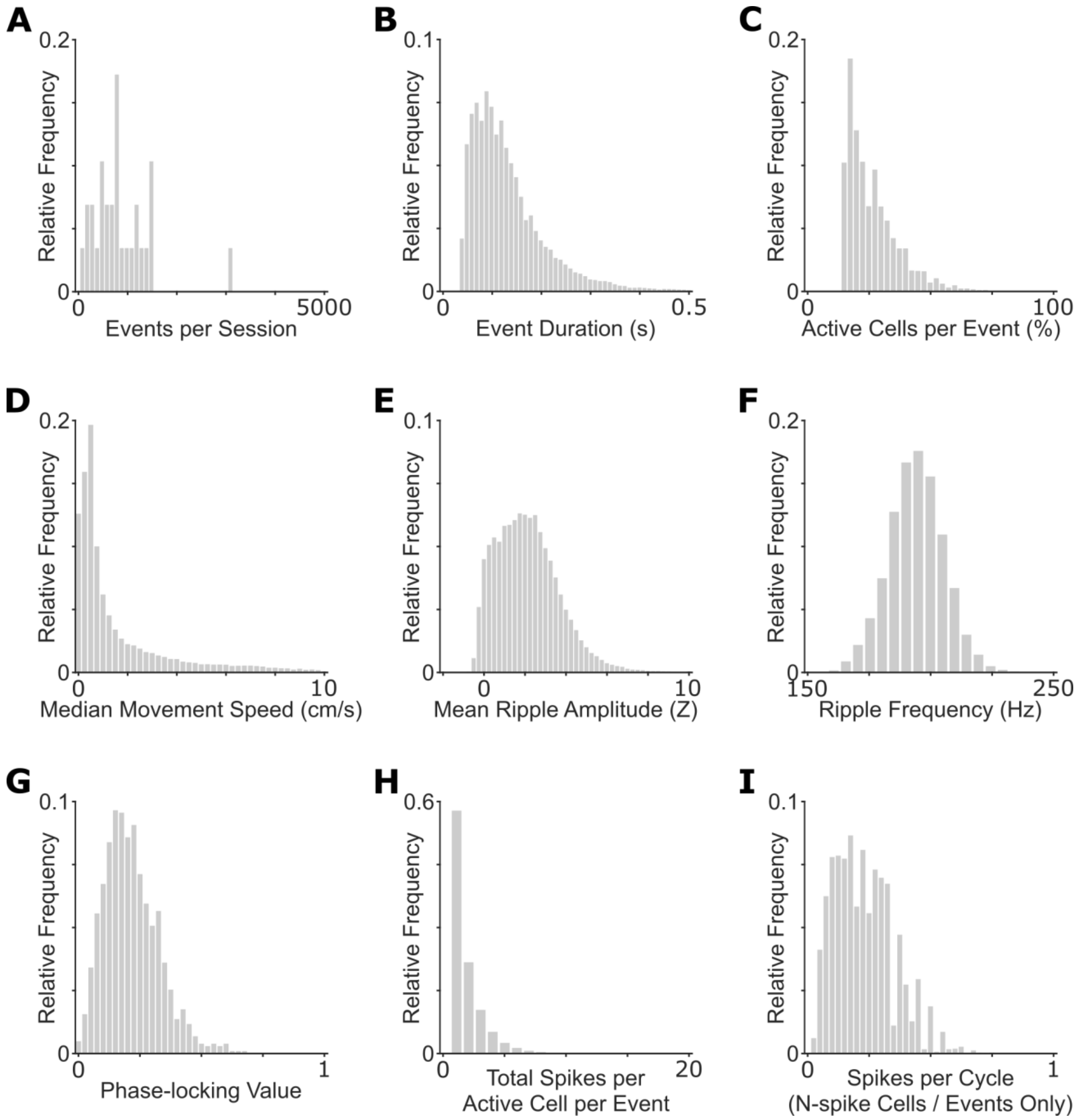


**Figure S1:** Details of CA1 candidate replay events detected using MUA. **[A]** Events per session (median±SD=759±599); **[B]** Event duration (114±71.3ms); **[C]** Active cells per event (23.3±10.2%); **[D]** Median within-event movement speed (0.661±2.0cm/s); **[E]** Mean within-event ripple power (Z=1.98±1.52); **[F]** Mean within-event ripple frequency (194±11.6Hz); **[G]** Ripple band phase locking across cells (0.199±0.112, significant phase locking in 69.4% of cells overall); **[H]** Spikes fired within-event per active cell (1±1.4); **[I]** Spikes fired per cycle by n-spike cells during n-spike events (0.179±0.127)

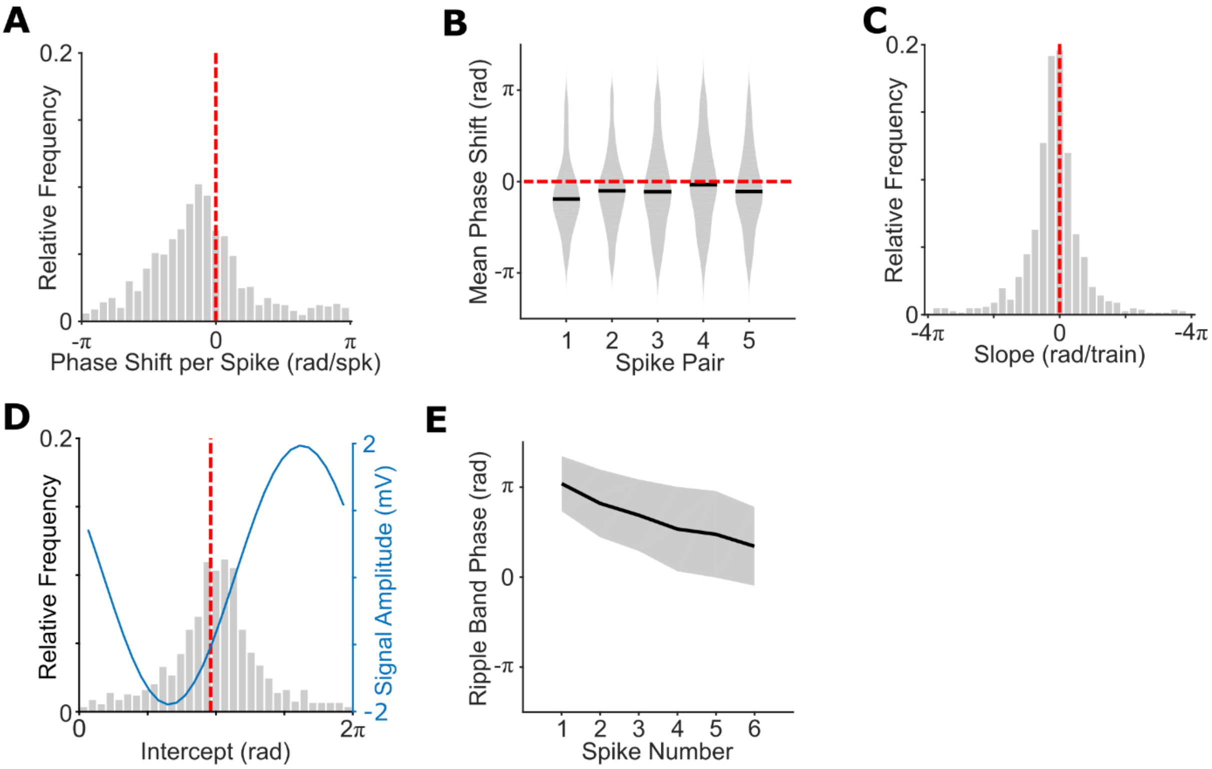


**Figure S2:** Changes in place cell ripple band firing phase during CA1 candidate replay events detected using MUA, when only the first spike fired in each oscillatory cycle is included. **[A]** Circular mean phase shift between all successive spikes in candidate replay events for each cell (n=951, overall circular median±circular SD=-0.553±1.13 rad/spk). This distribution is non-uniform (Rayleigh test, z=265, p<0.001) with a median value that differs significantly from zero (circular median test, p<0.001); **[B]** Circular mean phase shift by within-event spike pair, averaged across all cells. Each phase shift is non-uniformly distributed (all p<0.001) with a median value that differs significantly from zero (p<0.001) for all except the fourth spike pair (p=0.06); **[C]** Distribution of normalised time within spike train vs. ripple band phase slopes across cells (overall median±SD=-0.459±2.74 rad), which differs significantly from zero (t(947)=-5.91, p<0.001); **[D]** Distribution of normalised time within spike train vs. ripple band phase intercepts across cells (overall circular mean±circular SD=3.14±1.0 rad). This distribution is non-uniform (Rayleigh test, z=348, p<0.001) with a median value that differs significantly from the preferred firing phase of each cell (3.01±0.79 rad marked with a red dashed line, circular median test, p<0.001); **[E]** Unwrapped ripple band phase by spike number within candidate replay events, averaged across all cells (error bars indicate circular standard deviation)

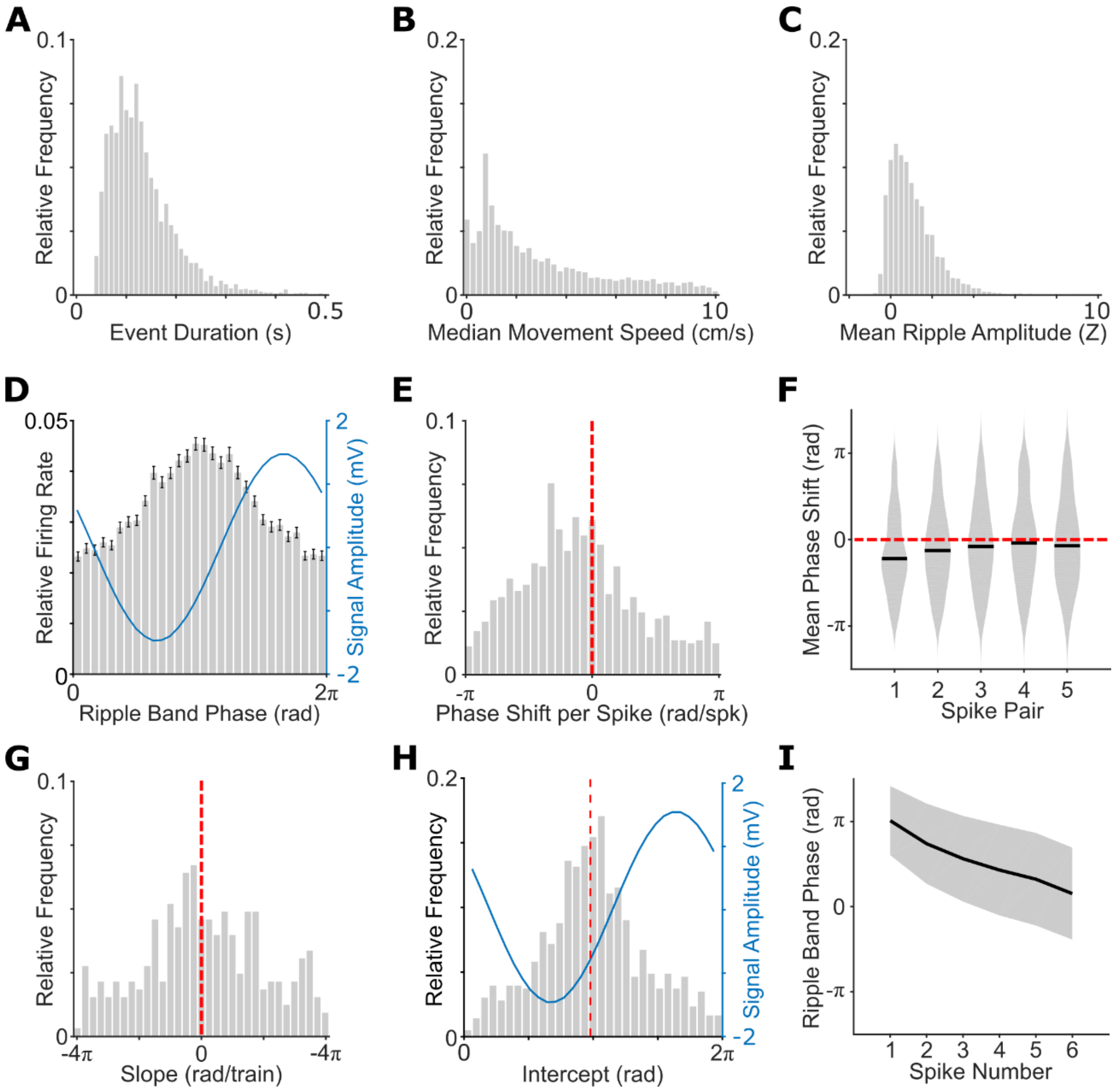


**Figure S3**: Changes in place cell ripple band firing phase during online CA1 candidate replay events detected using MUA. Total of 5590 events included (median±SD=133±146 per session, range 62-593), with 25±9.91% cells active per event, firing 1±1.56 spikes in total. **[A]** Event duration (117±64.4ms); **[B]** Median within-event movement speed (1.92±2.5cm/s); **[C]** Mean within-event ripple band power (Z=0.12±0.167); **[D]** Relative firing rate and mean LFP amplitude by ripple band phase, with significant phase locking in 26.9% of cells overall; **[E]** Circular mean phase shift between all successive spikes in candidate replay events (overall circular mean±circular SD=-0.649±1.51 rad/spk). This distribution is non-uniform (Rayleigh test, z=84.3, p<0.001) with a median value that differs significantly from zero (circular median test, p<0.001); **[F]** Circular mean phase shift by within-event spike pair, each of which are non-uniformly distributed (all p<0.001) with a median value that differs significantly from zero (all p<0.01) for all except the fourth and fifth spike pairs (both p>0.3); **[G]** Distribution of mean normalised time vs. ripple band phase slopes across cells (overall median±SD=-0.299±3.80 rad), which differs significantly from zero (t(864)=-2.23, p<0.05); **[H]** Distribution of mean normalised time vs. ripple band phase intercepts (overall circular mean±circular SD=3.09±1.31 rad). These are non-uniformly distributed (Rayleigh test, z=155, p<0.001), but their median value does not differ significantly from the preferred firing phase of each cell (3.08±1.08 rad, marked with a red dashed line); **[I]** Unwrapped ripple band phase by spike number within candidate replay events, averaged across all cells (error bars indicate circular standard deviation)

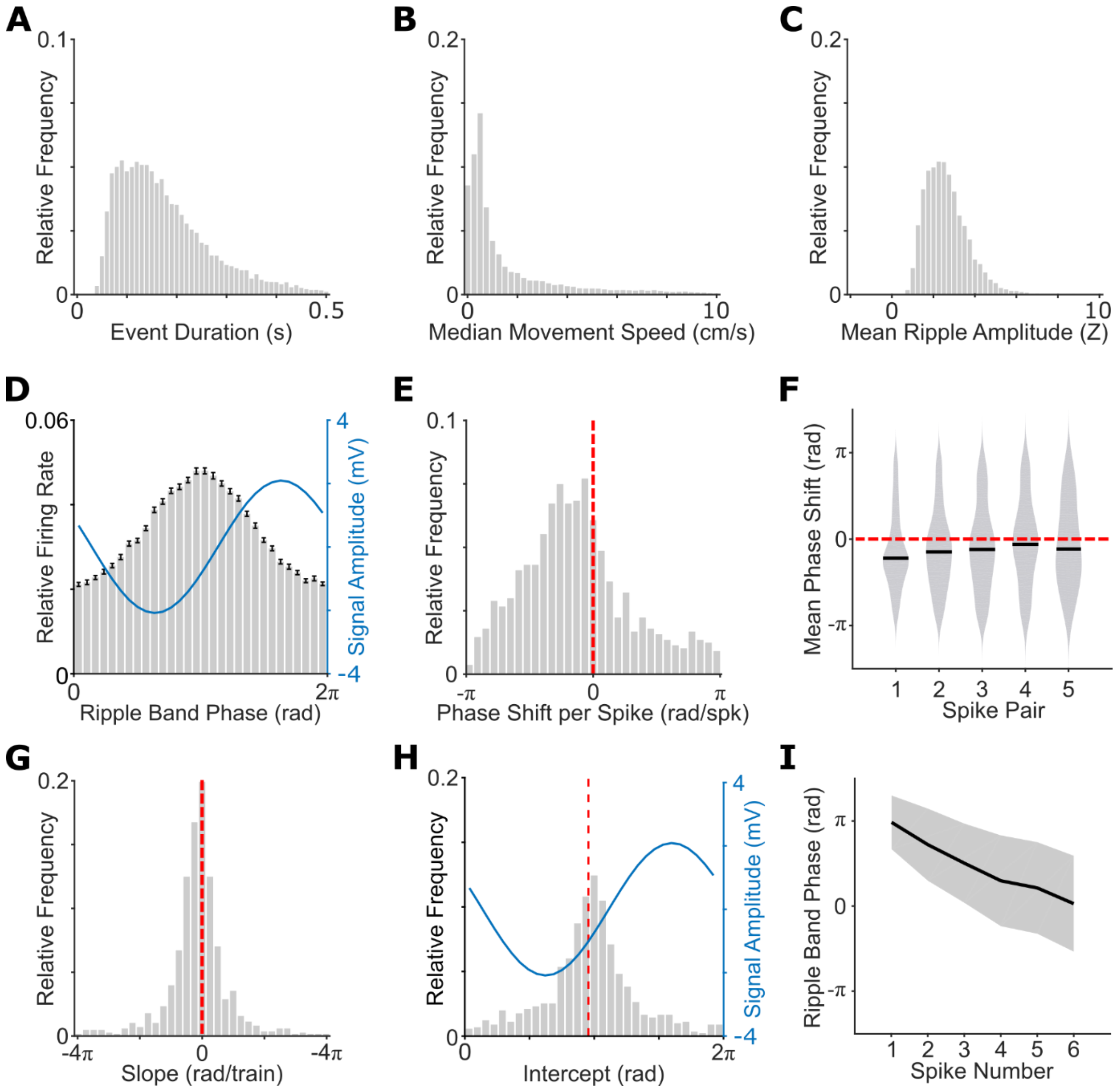


**Figure S4**: Changes in place cell ripple band firing phase during offline CA1 candidate replay events detected using increased ripple band power. Total of 20151 events included (median±SD=686±411 per session, range 112-2175), with 25.0±10.6% cells active per event, firing 1±1.31 spikes in total. **[A]** Event duration (155±90.2ms); **[B]** Median within-event movement speed (0.617±1.86cm/s); **[C]** Mean within-event ripple band power (Z=2.47±0.982); **[D]** Relative firing rate and mean LFP amplitude by ripple band phase, with significant phase locking in 70.3% of cells overall; **[E]** Circular mean phase shift between all successive spikes in candidate replay events (overall circular mean±circular SD=-0.653±1.33 rad/spk). This distribution is non-uniform (Rayleigh test, z=158.8, p<0.001) with a median value that differs significantly from zero (circular median test, p<0.001); **[F]** Circular mean phase shift by within-event spike pair, each of which are non-uniformly distributed (all p<0.001) with a median value that differs significantly from zero (all p<0.05); **[G]** Distribution of mean normalised time vs. ripple band phase slopes across cells (overall median±SD=-0.354±2.91 rad), which differs significantly from zero (t(916)=-4.45, p<0.001); **[H]** Distribution of mean normalised time vs. ripple band phase intercepts (overall circular mean±circular SD=3.03±1.05 rad). These are non-uniformly distributed (Rayleigh test, z=303, p<0.001), but their median value does not differ significantly from the preferred firing phase of each cell (3.0±0.783 rad, marked with a red dashed line); **[I]** Unwrapped ripple band phase by spike number within candidate replay events, averaged across all cells (error bars indicate circular standard deviation)

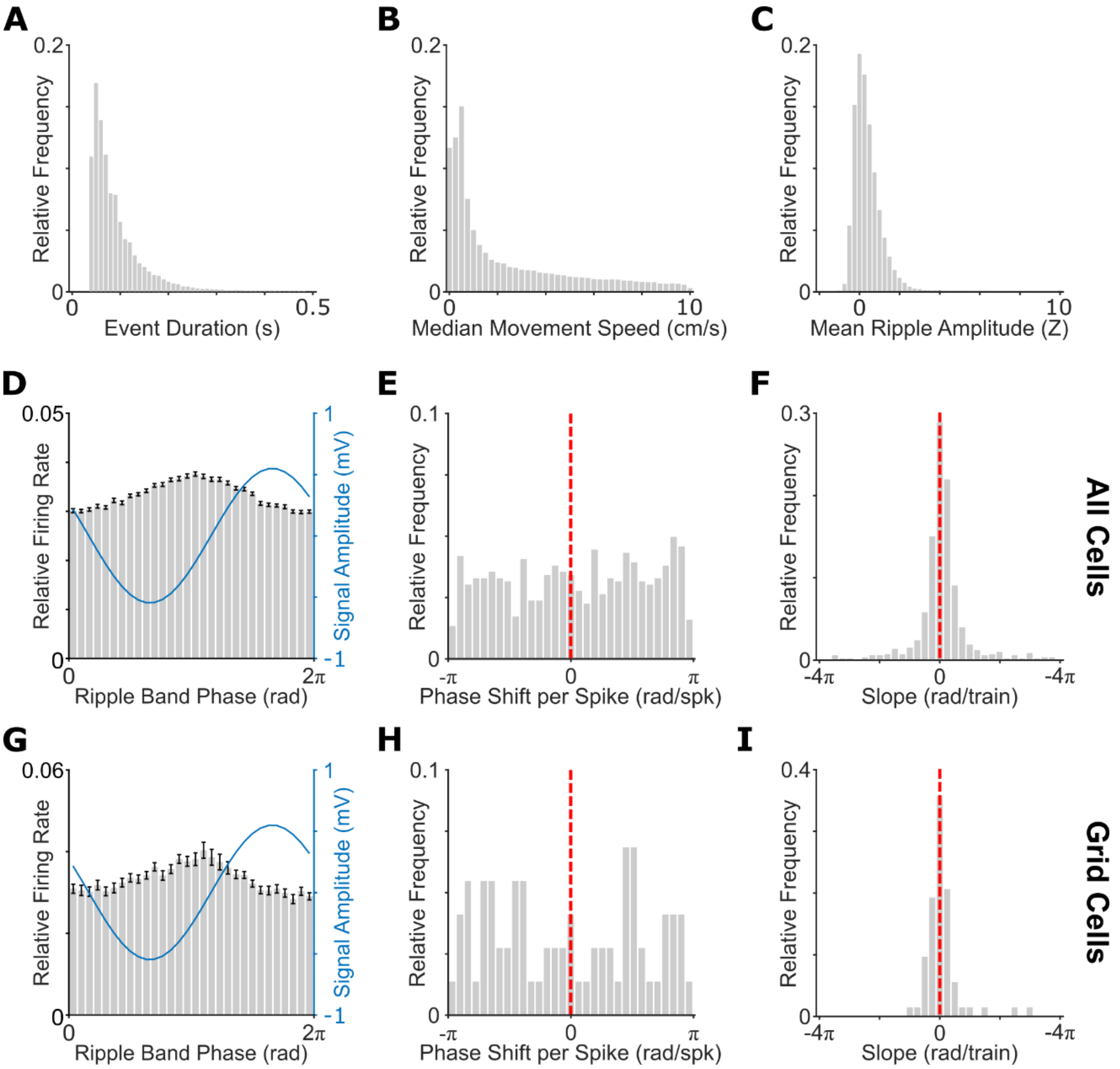


**Figure S5**: Changes in the ripple band firing phase of putative principal cells during offline MEC candidate replay events detected using MUA. We recorded a total of 877 cells in the deeper layers of MEC during REST, of which 832 were classed as putative principal cells (median±SD=31±9.94 per session, range 12-46). Next, we identified a total of 67540 candidate replay events (2701±1144 per session, range 268-4743), with 25±9.96% cells active per event firing 1±1.44 spikes in total. **[A]** Event duration (73±54.7ms); **[B]** Median within-event movement speed (1.03±2.61cm/s); **[C]** Median within-event ripple band power (Z=0.253±0.75); **[D]** Preferred ripple band firing phase across cells, with phase locking values of 0.07±0.08 across cells and significant phase locking in 42.1% of cells overall; **[E]** Circular mean phase shift between all successive spikes in each candidate replay event across 769/832 n-spike cells. This distribution is non-uniform (Rayleigh test, z=3.02, p=0.05) with a median value that does not differ significantly from zero (circular median test, p=0.15), but does differ significantly from the phase shift across CA1 cells (Watson-Williams test, F=215, p<0.001); **[F]** Distribution of mean normalised time vs. ripple band phase slopes across cells (overall median±SD=0.172±2.52 rad), which is significantly positive across cells (t(768)=2.63, p<0.01) and differs significantly from mean slopes across CA1 cells (t(1737)=-6.44, p<0.001); **[G]** Preferred ripple band firing phase across 73/832 n-spike grid cells, with phase locking values of 0.066±0.12 across grid cells and significant phase locking in 46.7% of those cells overall; **[H]** Circular mean phase shift between all successive spikes in each candidate replay event across n-spike grid cells. This distribution is uniform (Rayleigh test, z=0.848, p=0.43) and differs significantly from the phase shift across CA1 cells (Watson-Williams test, F=26.5, p<0.001); **[I]** Distribution of mean normalised time vs. ripple band phase slopes (overall median±SD=-0.119±1.83 rad), which is not different from zero across n-spike grid cells (t(72)=1.13, p=0.26) but does differ significantly from mean slopes across CA1 cells (t(1042)=-2.66, p<0.01)

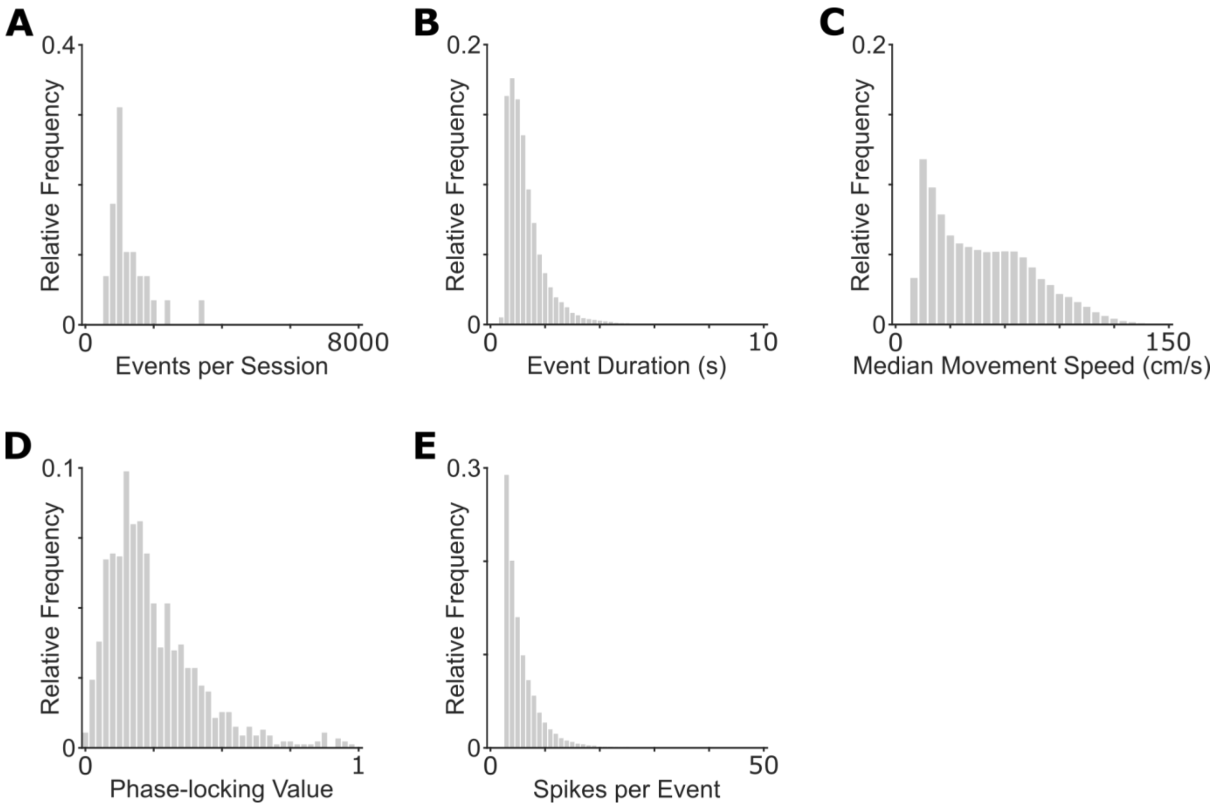


**Figure S6**: Details of CA1 theta spike trains, when only the first spike fired in each oscillatory cycle is analysed. Total of 37263 events included. **[A]** Events per session (median±SD=1080±602 per session, range 565-3466); **[B]** Event duration (1.1±0.70s); **[C]** Median within-event movement speed (42.4±27.4cm/s); **[D]** Theta phase locking across cells (0.199±0.165, significant phase locking in 37.1% of cells overall); **[E]** Spikes fired within each event (5±3.17)

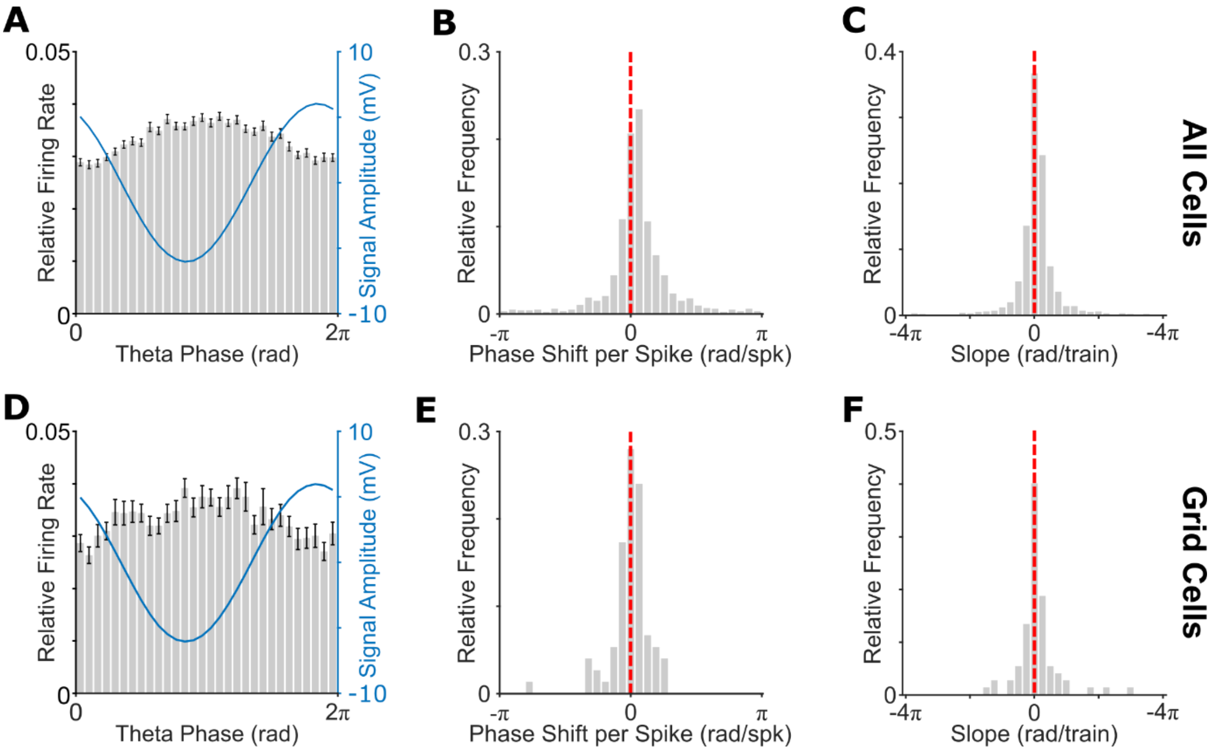


**Figure S7**: Changes in MEC putative principal cell theta firing phase during n-spike trains. We recorded a total of 1112 cells in the deeper layers of MEC during RUN, of which 1033 were classed as putative principal cells (median±SD=37±13.4 per session, range 15-82). Next, we identified a total of 121538 theta n-spike trains (4635±1936 per session, range 856-7218), lasting 1.53±1.37 s and incorporating 5.0±6.66 spikes per train. **[A]** Relative firing rate and mean signal amplitude by theta band phase, with median±SD phase locking values of 0.188±0.164 across cells and significant phase locking in 59.4% of cells; **[B]** Circular mean phase shift between all successive spikes in theta n-spike trains across 1011/1033 n-spike cells (overall circular mean±circular SD=0.171±0.714 rad/spk). This distribution is non-uniform (Rayleigh test, z=607, p<0.001) with a median value that differs significantly from zero (circular median test, p<0.001); **[C]** Distribution of normalised within n-spike train time vs. theta phase slopes (overall median±SD=0.185±1.74 rad), with a mean value which differs significantly from zero (t(1008)=3.99, p<0.001); **[D]** Relative firing rate and mean signal amplitude by theta band phase across 75/1033 n-spike grid cells, with phase locking values of 0.189±0.156 across grid cells and significant phase locking in 63.9% of those cells; **[E]** Circular mean phase shift between successive spikes in theta n-spike trains of grid cells (overall circular mean±circular SD=0.0133±0.457 rad/spk). This distribution is non-uniform (Rayleigh test, z=60.8, p<0.001) with a median value that does not differ significantly from zero (circular median test, p=0.49); **[F]** Distribution of normalised within n-spike train time vs. theta phase slopes (overall median±SD=0.119±2.02 rad), with a mean value which does not differ significantly from zero across grid cells (t(74)=1.08, p=0.29)

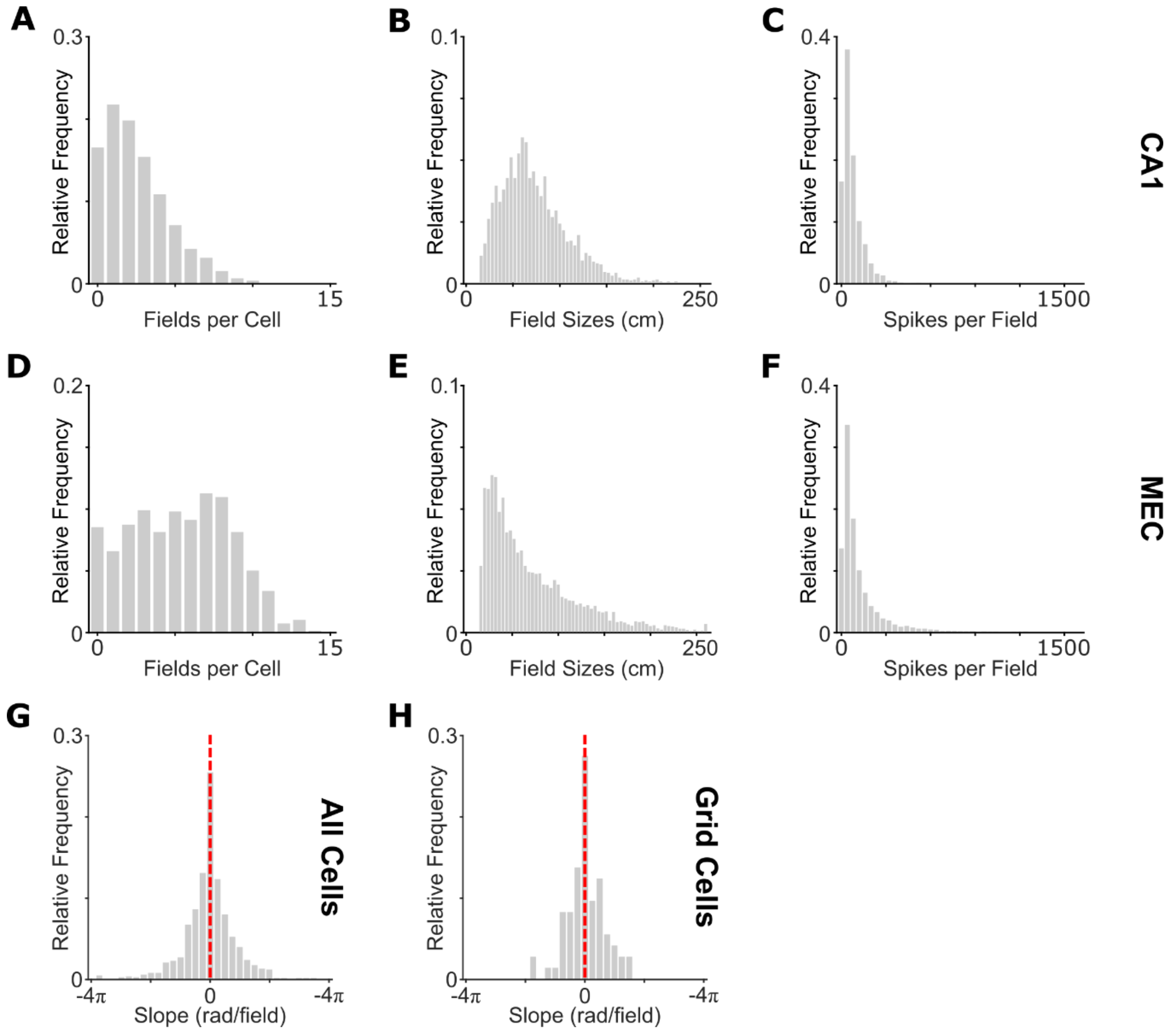


**Figure S8:** Spatial firing field properties in CA1 and MEC, and the relationship between actual location and theta firing phase in MEC. **[A]** CA1 spatial firing fields per cell, summed across inbound and outbound rate maps (median±SD=2±2.17, range 0 to 10); **[B]** CA1 firing field sizes (68±35.7cm, range 20 to 254cm); **[C]** Total spikes fired over all movement trajectories through each firing field (54±65.3, range 5 to 540); **[D]** MEC spatial firing fields per cell, summed across inbound and outbound rate maps (6±3.3, range 0 to 14). 946/1033 putative principal cells (91.6%) active during RUN have ≥1 field included on either outbound or inbound runs along the track; **[E]** MEC firing field sizes (58±49.1cm, range 20 to 316cm); **[F]** Total spikes fired over all movement trajectories through each firing field (65±128, range 5 to 1251); **[G]** Distribution of mean normalised location vs. theta phase slopes across MEC principal cells (-0.00628±2.56 rad/field), which does not differ significantly from zero ($t(944)=-0.753$, $p=0.452$); **[H]** Distribution of mean normalised location vs. theta phase slopes (0.0792±2.03 rad/field) across 73/1033 grid cells that passed our criteria for inclusion, which does not differ significantly from zero ($t(72)=0.344$, $p=0.732$). In total, 78 / 536 grid fields (14.6%) showed a significant negative correlation between location within the firing field and theta phase, which is more than expected by chance (binomial test, $p<0.001$)

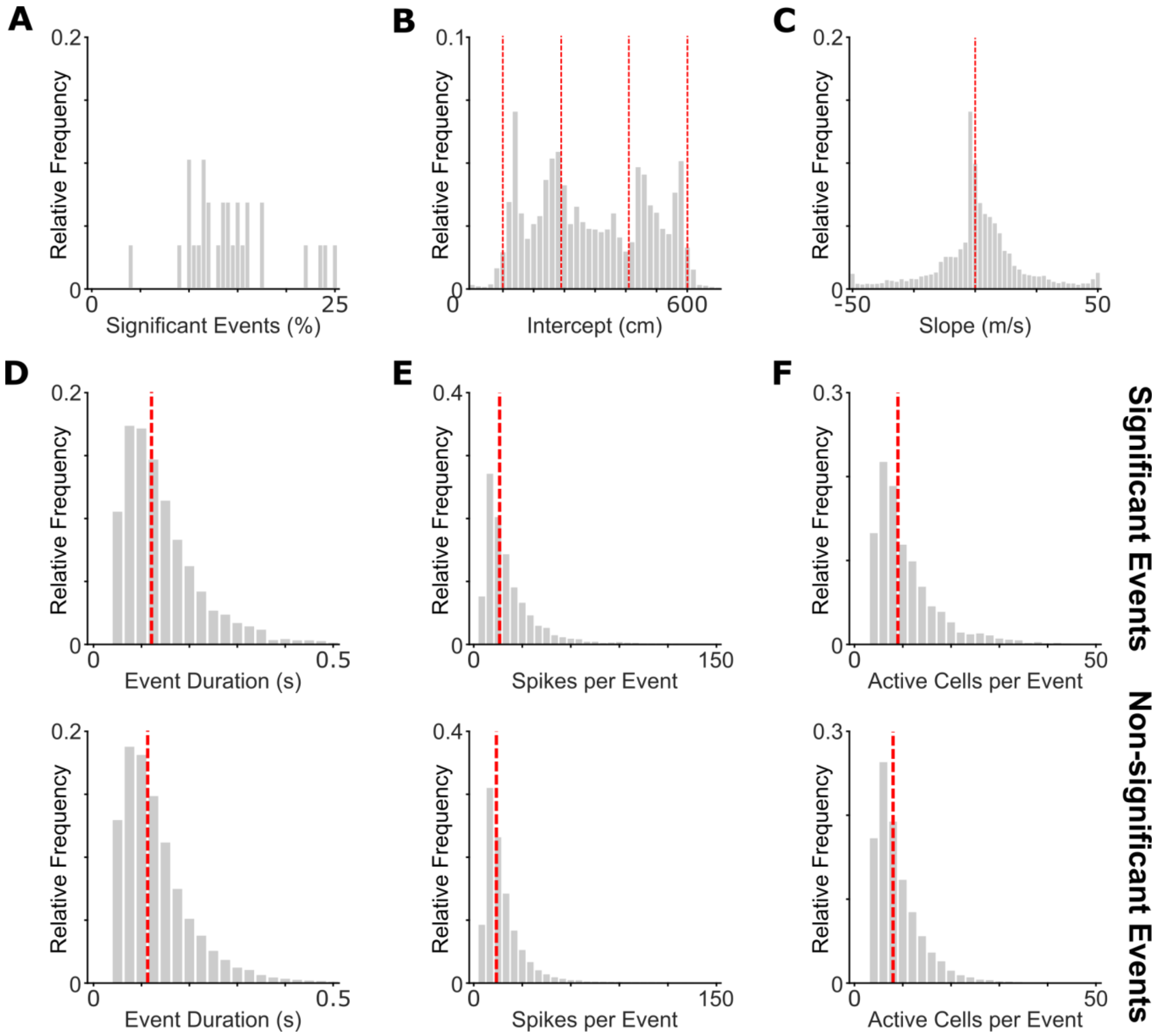


**Figure S9:** Details of significant CA1 replay trajectories during REST. **[A]** Proportion of significant events per session (median±SD=13.5±4.7%, range 4.2-24.9%); **[B]** Intercept (i.e. origin) of significant trajectories. Note that these tend to cluster around the ends and corners of the Z-track (marked with red dashed lines); **[C]** Speed of significant trajectories (absolute median±SD=7.5±12.6m/s). Positive values (57.7% of all events) correspond to forward and negative values (42.3% of all events) to reverse replay events. There is a significant bias towards forward replay (binomial test, p<0.001); **[D]** Significant events (upper panel, median=121ms) are longer than non-significant events (lower panel, median=113ms, Mann-Whitney U-test, Z=7.31, p<0.001); **[E]** Significant events (upper panel, median=16 spikes) incorporate more spikes than non-significant events (lower panel, median=14 spikes, Mann-Whitney U-test, Z=11.6, p<0.001); **[F]** Significant events (upper panel, median=9 active cells) incorporate more active cells than non-significant events (lower panel, median=8 active cells, Mann-Whitney U-test, Z=13.8, p<0.001)

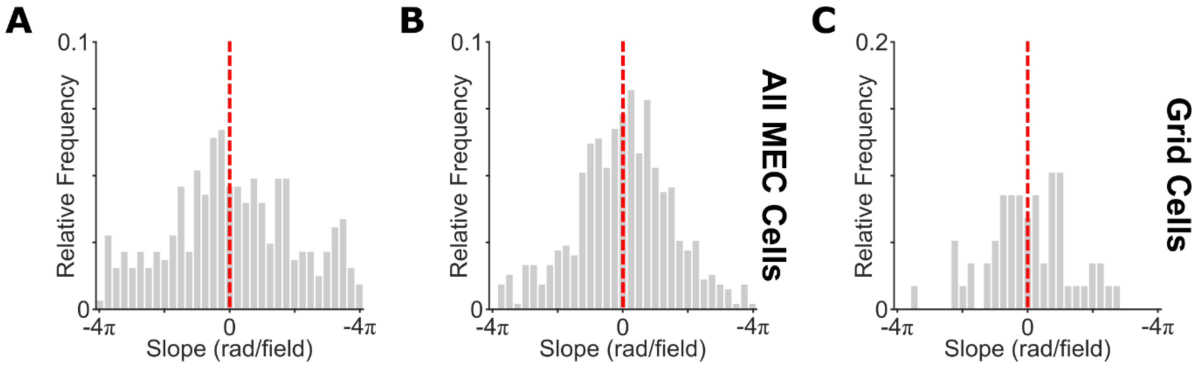


**Figure S10**: Relationship between decoded location and ripple band firing phase during online replay events, and in MEC. **[A]** In CA1, a total of 1177/5590 online replay events corresponded to significant trajectories along the track (median±SD=20±9.16% events per session). A total of 721 place fields subsequently passed our threshold for inclusion (≥5 spikes covering ≥50% of the firing field), with 360/1044 cells (34.5%) having ≥1 field included and 12±12.3 spikes per field (range 5 to 115). Distribution of mean decoded location vs. ripple phase slopes across CA1 place cells (-0.289±6.97 rad/field), which does not differ significantly from zero (t(359)=0.067, p=0.95); **[B]** In MEC, a total of 7211/67540 offline replay events corresponded to significant trajectories along the track (median±SD=10.9±3.11% events per session). A total of 2442 firing fields passed our threshold for inclusion (≥5 spikes covering ≥50% of the firing field), with 572/877 cells (65.2%) having ≥1 field included and 16±37.4 spikes per field (range 5 to 602). Distribution of mean decoded location vs. ripple phase slopes across MEC principal cells (0.299±4.65 rad/field), which does not differ significantly from zero (t(550)=0.326, p=0.744); **[C]** Distribution of mean decoded location vs. ripple phase slopes (-0.00628±4.02 rad/field) across 59/877 grid cells that passed our criteria for inclusion, which does not differ significantly from zero (t(58)=0.310, p=0.758)
